# Sex-specific topology of the nociceptive circuit shapes dimorphic behavior in *C. elegans*

**DOI:** 10.1101/2021.12.14.472335

**Authors:** Vladyslava Pechuk, Yehuda Salzberg, Gal Goldman, Aditi H. Chaubey, R. Aaron Bola, Jonathon R. Hoffman, Morgan L. Endreson, Renee M. Miller, Noah J. Reger, Douglas S. Portman, Denise M. Ferkey, Elad Schneidman, Meital Oren-Suissa

**Author notes:** These authors contributed equally (Vladyslava Pechuk, Yehuda Salzberg, Gal Goldman). Correspondence |.

## Abstract

How sexually dimorphic behavior is encoded in the nervous system is poorly understood. Here, we characterize the dimorphic nociceptive behavior in *C. elegans* and study the underlying circuits, which are composed of the same neurons but are wired differently. We show that while sensory transduction is similar in the two sexes, the downstream network topology markedly shapes behavior. We fit a network model that replicates the observed dimorphic behavior in response to external stimuli, and use it to predict simple network rewirings that would switch the behavior between the sexes. We then show experimentally that these subtle synaptic rewirings indeed flip behavior. Strikingly, when presented with aversive cues, rewired males were compromised in finding mating partners, suggesting that network topologies that enable efficient avoidance of noxious cues have a reproductive “cost”. Our results present a deconstruction of the design of a neural circuit that controls sexual behavior, and how to reprogram it.

## INTRODUCTION

Sexual identity is an obvious source of variation in phenotypic traits. In sexually reproducing species, for example, females and males often respond to sensory cues in sexually dimorphic ways, reflected in their foraging, courtship, mating, and aggressive behaviors (Hashikawa et al., 2018; McKinsey et al., 2018). Such behavioral differences may originate from multiple sources or their combination: (1) Sex-specific neuronal populations, such as the courtship-related male-specific P1 neuronal cluster in Drosophila (Kimura et al., 2008) or the egg-laying related hermaphrodite-specific cholinergic VC and serotonergic HSN neurons in *C. elegans* (White et al., 1986). (2) Differences in the response properties of sensory neurons, such as in auditory tuning in frogs (Keddy-Hector et al., 1992; McClelland et al., 1997; Narins and Capranica, 1976; Shen et al., 2011; Wilczynski et al., 1984), sex-limited expression levels of G-protein-coupled receptors (GPCRs) in the AWA sensory neuron in *C. elegans* (Ryan et al., 2014; Wan et al., 2019), or the activation of the mouse vomeronasal organ (VNO) in response to male and female urine (He et al., 2008; Holy et al., 2000). (3) Different topologies of neural circuits over the same neurons, as has been reported in *C. elegans* (Cook et al., 2019) and in the pre-optic area of rats (Raisman and Field, 1971, 1973). In mice, complex social behaviors are shaped by adaptive modulatory changes, although it is unclear whether these behaviors are a result of dimorphic connectivity or of dimorphic gene expression (Bayless and Shah, 2016; Cooke and Woolley, 2005; Li and Dulac, 2018). (4) Differences in synaptic strengths over the same topology (Tikhonov and Bialek, 2016). (5) Different circuit functions due to neuromodulation (Jang et al., 2012).

To delineate the role and impact of these potential sources in shaping dimorphic behaviors, we here explore the universal trait of the noxious stimulus response. *C. elegans* presents us with a unique opportunity to disentangle the design and function of neuronal circuits that generate dimorphic behaviors, since differences in topology have been mapped for both sexes (Cook et al., 2019; Jarrell et al., 2012; White et al., 1986), the nervous system is accessible at the resolution of single identified neurons and connections, and the behavioral repertoire is well characterized (Stephens et al., 2008). Moreover, the small size of functional sub-networks allows circuit design to be explored by a detailed computational model.

Pain tolerance and dimorphic responses in males and females have been well documented (McCarthy, 2021; Mogil, 2012, 2020; Sorge and Strath, 2018; Wiesenfeld-Hallin, 2005), whereas the underlying neural mechanisms have only recently been addressed (Sorge et al., 2015; Yu et al., 2021). In *C. elegans*, many noxious cues are sensed and transduced by a single pair of glutamatergic neurons, the ASHs (Bargmann et al., 1990)(Figure 1A), and the downstream circuit is compact. We, therefore, explored the relationship between sex-specific network topology and the resulting behavioral output by characterizing sensory transduction and differences between the processing circuits, and manipulating the circuit *in silico* and *in vivo*.

**Figure 1.**
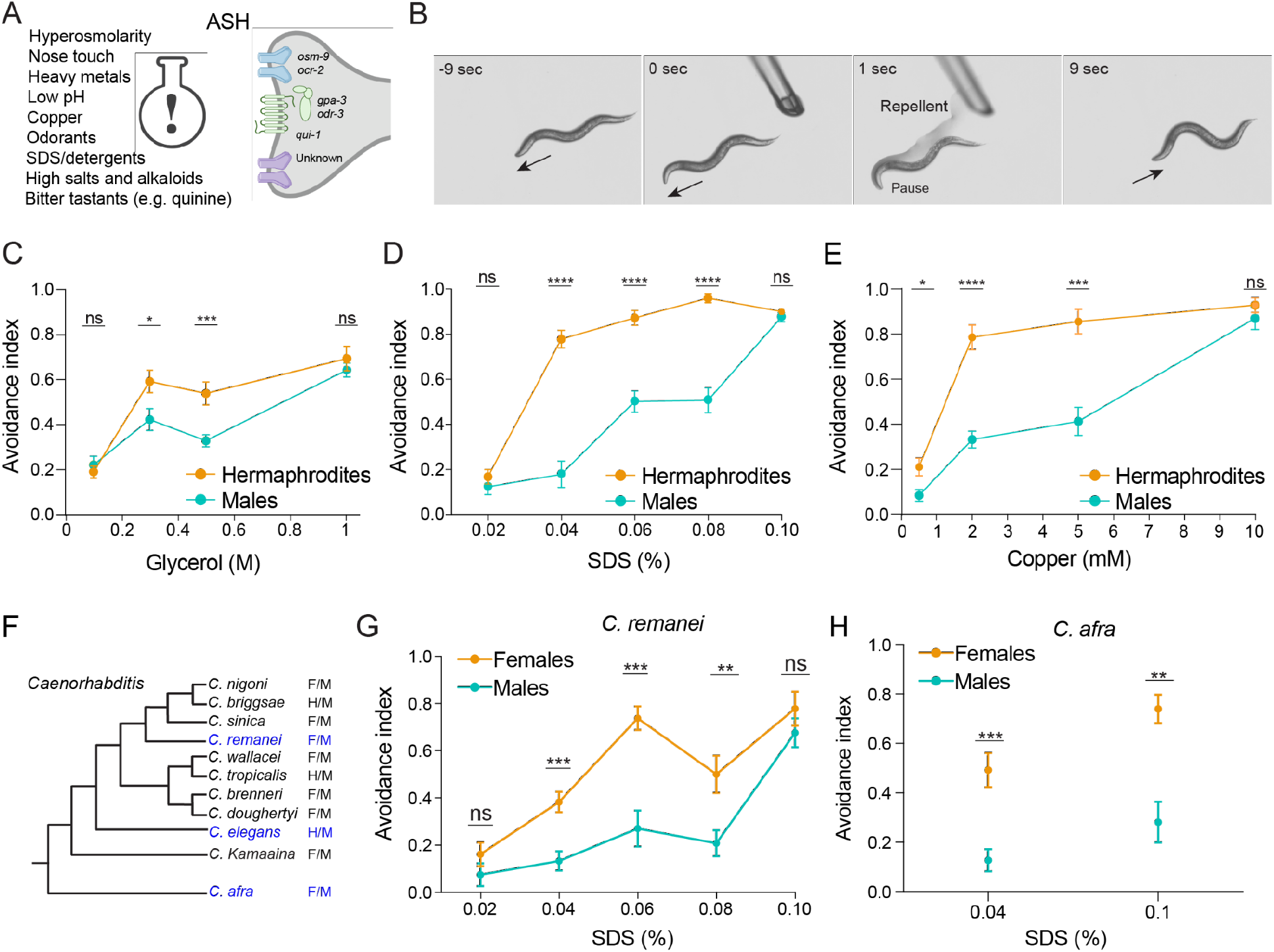
*C. elegans* exhibits sexually dimorphic behaviors in response to nociceptive stimuli. (A) ASH is a polymodal nociceptive neuron that senses aversive cues through a variety of receptors and signaling molecules. (B) Tail drop test: As the animal moves forward on an agar plate, (−9 sec), a glass micropipette containing a repellent delivers a drop near the tail that reaches the anterior extremity of the animal by capillary action (1 sec). This causes the animal to pause and initiate the avoidance response by moving backward (9 sec). (C-E) Sexually dimorphic dosedependent behavioral responses to aversive stimuli: (C) glycerol, (D) SDS, and (E) copper (tail drop assay, see Methods for a full description of behavioral tests). The avoidance index represents an average of the fraction of reversal responses (scored as reversing or not reversing) in 10 assays of each single animal. n = 9-15 worms per group. (F) Cladogram of the *Caenorhabditis* genus, where two alternative mating strategies exist. F=female, M=male, H=hermaphrodite. (G-H) The sexually dimorphic behavioral responses are evolutionarily conserved also in the female-male species *C. remanei* (G) and *C. afra* (H) (tail drop assay). n = 8-13 worms per group. Error bars are standard error of the mean (SEM). We performed a Mann-Whitney test for all comparisons, **** *p* < 0.0001, *** *p* < 0.001, ** *p* < 0.01, * *p* < 0.05, ns - non-significant.

We show that sensory transduction is similar between the two sexes and that differences in the topology of the circuit that processes the nociceptive information are sufficient to explain the observed behavioral differences. By simulating the dimorphic networks, we found a small range of circuit and neuronal parameters for which the behavioral output of the simulation matched the worms’ experimentally defined behavior. We then used our model to identify critical rewirings of the hermaphrodite and male circuits that might reshape behavior. We validated these predictions experimentally, and showed that while the hermaphrodite circuit is robust to perturbations, the male circuit could be manipulated to generate the responses of the opposite sex - even by changing a single connection. Moreover, males with feminized avoidance behavior were compromised in finding mates, when simultaneously presented with aversive cues. Our results thus suggest that sexual identity sculpts neuronal circuits for the sex-specific needs of the organism, even for traits necessary for one’s survival.

## RESULTS

### *C. elegans* displays sexually dimorphic behaviors in response to nociceptive stimuli

To explore the nociceptive responses of the two sexes, we presented *C. elegans* hermaphrodites and males with aversive stimuli and measured their tail drop and head drop assay responses (Figure 1B, see Methods). We found that the avoidance responses of the two sexes differed for diverse types of noxious stimuli: SDS (chemical avoidance), glycerol (high osmolarity), quinine (alkaloid) and copper (heavy metal) (Figure 1C-E, Figure S1A-D). Overall, hermaphrodites showed a stronger dose-dependent avoidance behavior compared to males. This observation remained valid even for gonochoric (female-male, as opposed to hermaphrodite-male) species of the *Caenorhabditis* genus, such as *C. remanei* and *C. afra* (Figure 1F-H, S1E), suggesting that the dimorphic chemo-avoidance behavior is strongly conserved among nematoda, regardless of the reproductive mode.

### Similar activation of sensory neurons induces dimorphic avoidance behaviors

Having established nociceptive behavioral dimorphism, we turned to explore the underlying neural circuits in the two sexes (Figure 2A): located downstream of the key sensory neuron ASH are the interneurons AVA and AVD, which mediate backward movement, and the interneurons AVB and PVC, which mediate forward movement (Gjorgjieva et al., 2014; Kawano et al., 2011; Patil et al., 2015; Zhen and Samuel, 2015). Organized in pairs, these neurons are located symmetrically on the right and left sides of the worm. The difference in activation between downstream A-type and B-type motor neurons determines the direction of movement (Kawano et al., 2011). Notably, the nociceptive circuits in hermaphrodites and males consist of the same set of neurons, which are organized in sexually dimorphic topologies (Cook et al., 2019; Jarrell et al., 2012; White et al., 1986).

**Figure 2.**
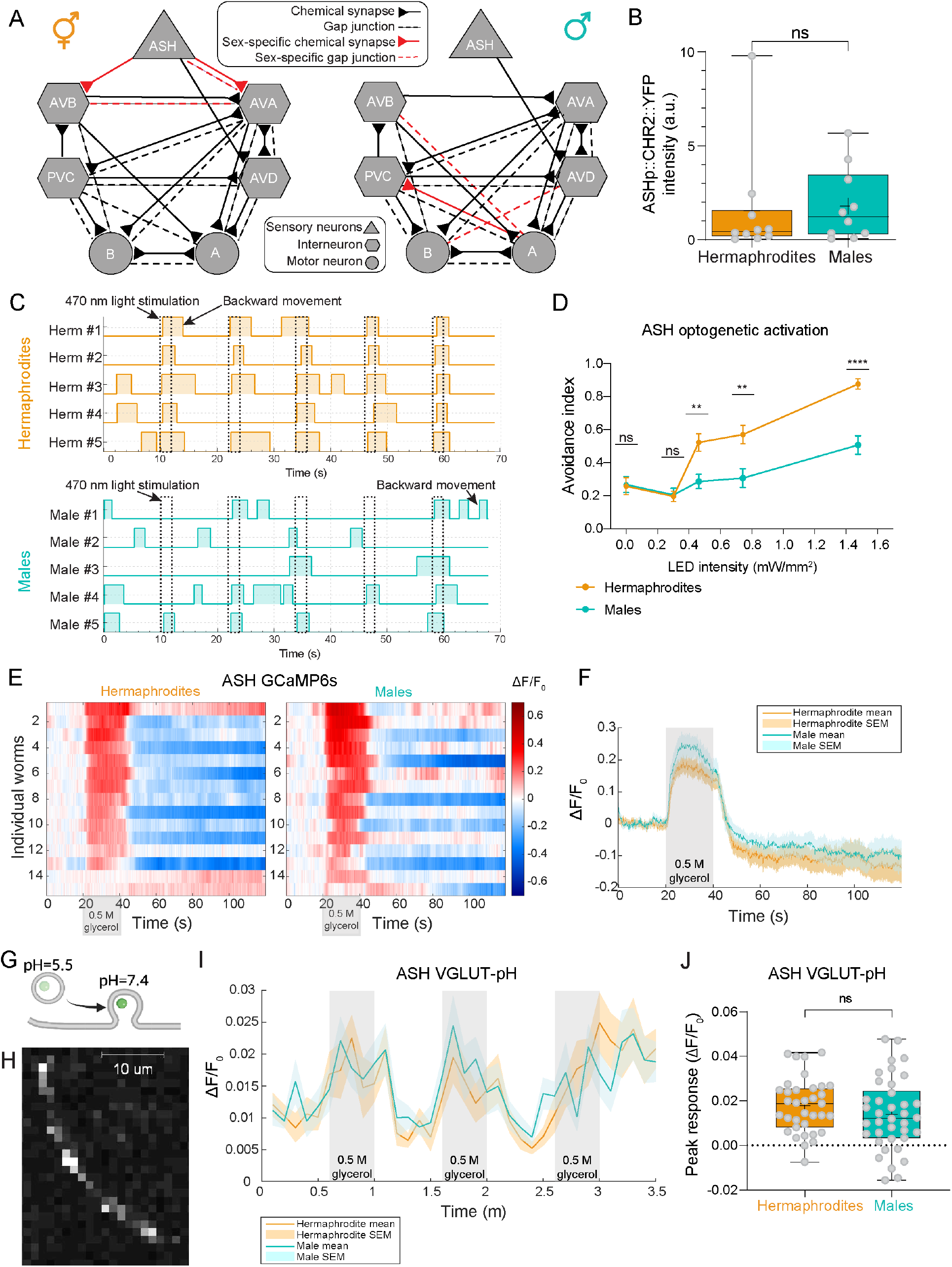
Sexually dimorphic behavioral choices are not encoded at the sensory level. (A) Hermaphrodite (left) vs. male (right) predicted connectivity (Cook et al., 2019; White et al., 1986). Chemical synapses are represented as arrows, gap junctions as dashed lines, and sexspecific connections as red arrows and dashed lines (for chemical or gap junction connections, respectively). (B) Quantification and comparison of ChR2 expression levels in ASH in both sexes, using the *ASHp::ChR2::YFP* transgene. Fluorescence intensity was measured with FIJI after confocal images were captured. n = 10 per sex. (C) Representative avoidance responses of five hermaphrodites and males to five consecutive ASH optogenetic activations (blue-light stimuli, marked with dashed lines). Plotted boxes represent reversal events. (D) Specific activation of ASH neurons with different LED intensities recapitulates the sexually dimorphic behavioral responses. n = 37-40 hermaphrodites, 30-40 males. See control groups without *all-trans-retinal* (ATR) in Figure S2A. (E) GCaMP6s calcium responses of ASH neurons to 0.5 M glycerol. Heatmaps represent the calcium levels of individual worms. The stimulus is applied at 20-40 seconds. Calcium levels are normalized and color-coded. (F) Average and SEM traces of ASH calcium responses of hermaphrodites and males. Gray background denotes the time of stimulus delivery. See Figure S2C for statistical comparison of peak responses. (G) Illustration of the function of the pHluorin reporter. The GFP signal increases upon vesicular exocytosis due to a change in the pH (Miesenböck et al., 1998). (H) Representative confocal micrograph of a ASH axon, with a visible fluorescent signal of *sra-6p::EAT-4::pHluorin*. Scale bar is 10 μm. (I) Average and SEM traces of glutamatergic secretion from ASH axons of the two sexes, in response to three stimuli (gray background). (J) Quantification of peak responses. Each dot is a single response, each worm has (at most) three dots in the graph. n = 35 hermaphrodite dots (12 animals), 38 male dots (14 animals). B and J are box and whiskers plots, vertical line represents the median and “+” is the mean. We performed a Mann-Whitney test for all comparisons, **** *p* < 0.0001, ** *p* < 0.01, ns - non-significant.

We first ascertained whether the dimorphic behavior can be reproduced by activating ASH directly. This entailed activating ASH optogenetically in both sexes using channelrhodopsin-2 (ChR2) and monitoring the responses (while validating that the ChR2 levels do not vary between the sexes; Figure 2B). Optogenetic activation of ASH was sufficient to elicit sexually dimorphic responses (Figure 2C). Moreover, a gradual increase in illumination intensities reproduced the dimorphic dose-dependent response curve of the worms, in agreement with the behavioral chemorepellent assays (Figure 2D, Figure S2A).

Next, we sought to determine whether ASH displays any dimorphism in intrinsic properties. We first measured the protein expression levels and subcellular localization of different receptors and signaling molecules in the two sexes. Since ASH is a polymodal sensory neuron (Bargmann et al., 1990; Bono and Maricq, 2005; Culotti and Russell, 1978; Ferkey et al., 2021; Kaplan and Horvitz, 1993; Troemel et al., 1995) we tested the protein expression levels of several ASH receptors and signal transduction molecules: OSM-9, OCR-2, OSM-10, GPA-3, QUI-1. All these proteins displayed non-dimorphic expression levels and subcellular localization (Figure S2B). We then imaged the activity of ASH neurons using transgenic worms expressing the genetically-encoded calcium indicator GCaMP6s. For the same glycerol concentration that elicits dimorphic behavioral responses (0.5 M glycerol, Figure 1C), the intracellular calcium response of the ASH sensory neurons was similar in the two sexes (Figure 2E-F, S2C). Recent work has shown that a pair of sex-shared sensory neurons in *C. elegans*, the PHCs, undergoes sexually dimorphic differentiation marked by sex-specific scaling of the synaptic vesicle machinery (Serrano-Saiz et al., 2017). We, therefore, measured the intensity of a fosmid-based reporter for the vesicular glutamate transporter *eat-4/VGLUT* in ASH and found that it was also non-dimorphic (Figure S2D-F). To monitor synaptic vesicle exocytosis and retrieval, we used the VGLUT-pHluorin sensor (Ventimiglia and Bargmann, 2017) to measure glutamate secretion from ASH axons following three repeated stimulations (Figure 2G-H). The delivery of 0.5 M glycerol to the head of the worms, where the ASH cilia are located, resulted in a non-dimorphic increase in VGLUT-pHluorin fluorescence (Figure 2I-J).

Thus, multiple lines of evidence pertaining to morphology, receptor and signal transduction, calcium response, vesicle packaging and neurotransmitter secretion suggest that sensory transduction is non-dimorphic. We, therefore, concluded that sexually-dimorphic avoidance behaviors likely result from differences occurring downstream of the sensory neurons.

### Simulating the circuit for nociception suggests a critical role for sex-specific connections in dimorphic behavior

Having established that the sensory inputs to the circuit are non-dimorphic, we turned to ask how the different topology of the networks of hermaphrodites and males (Figure 2A) shapes the dimorphic nociceptive behavior. Since experimental assessment and manipulation of all these differences simultaneously is challenging, we simulated the response of the two circuits to activation of the sensory neuron, and used the difference in activations between the motor neurons (A and B) to determine the predicted direction of movement (see Methods).

Since similar network activity may arise from very different sets of neuronal and network parameters (Prinz et al., 2004), and since the biophysical parameters of neurons in this circuit have not been measured simultaneously and in large populations – we explored a wide range of biologically plausible values for the circuit’s biophysical parameters (Goodman et al., 1998; Lindsay et al., 2011; Palyanov et al., 2012; Rakowski and Karbowski, 2017; Rakowski et al., 2013; Roehrig, 1998; Varshney et al., 2011; Wicks et al., 1996) (Figure 3A-B). For each of our 7 parameters, we used 7 different values; overall, we simulated 7^7^ = 823,543 different parameter combinations and compared the activity of the simulated motor neurons to the worms’ actual behavior in response to nociceptive stimuli. We found three distinct sets of parameter combinations (Figure 3C) for which the activity of the motor neurons in the simulation matched the worms’ experimentally measured behavior (Figure 1) in response to a strong sensory stimulus (which is expected to trigger high avoidance in both sexes): 1.9% of parameter combinations were appropriate only for the hermaphrodites, 1.2% were unique for males, and 0.3% were appropriate for both sexes (Figure 3C). The range of parameter values that we used was substantiated by the fact that most values were represented in at least one valid set (Figure 3D). The marginal distributions over the single parameters reflect the differences between the sexes, and that the values of the mutual sets may also differ considerably from those of the two sexes (Figure 3D; for example, valid parameter sets of hermaphrodites and males seem to favor different values of membrane resistance and synaptic conductance). Projecting the hermaphrodite-only, male-only, and mutual sets onto the 2 first principal components of the space of all valid sets (which capture ~45% of the variance) shows that the mutual set inhabits a compact part of the parameter space (compared to the male only and hermaphrodite only ones), whereas the differences between hermaphrodites and males are more intricate (Figure 3E).

**Figure 3.**
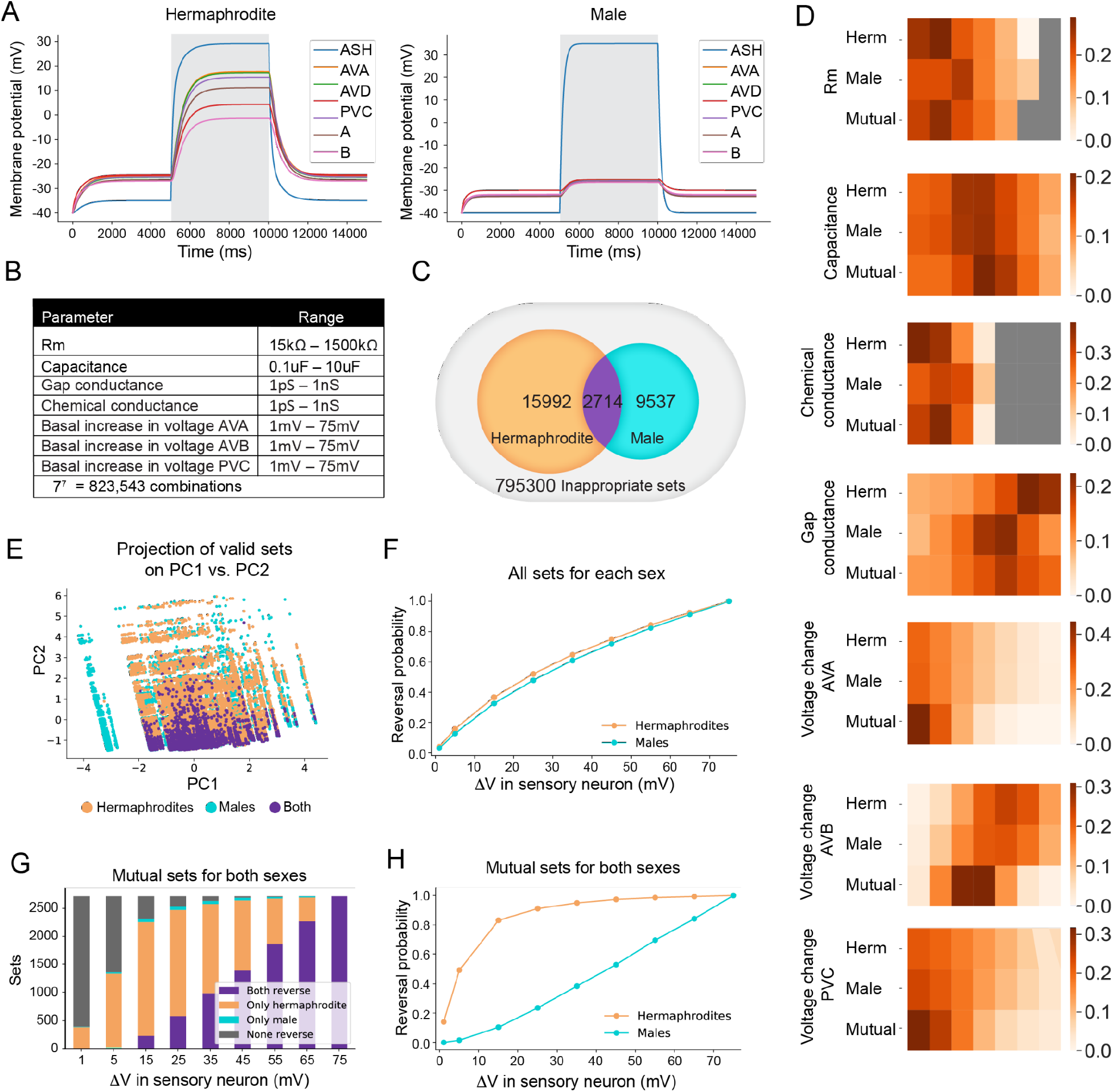
Simulation of the circuit for nociceptive behaviors recapitulates the behavioral differences between the sexes. (A) Representative simulation of the membrane potential of the cells in the circuit, for the hermaphrodite (left) and the male (right). The detection of an aversive stimulus was simulated using input current to the sensory neuron ASH for 5 seconds (5000 ms - 10000 ms, gray background). Constant basal input was given to the interneurons, such that it increased their membrane potentials in the following manner: AVA - 8.7 mV, PVC - 4.2 mV, AVB - 17.8 mV (see Methods for more details). Biophysical parameters that were used: Rm: 15 kΩ, capacitance: 10 uF, chemical conductance: 31.6 pS, gap conductance: 1 nS. All cells were simulated as excitatory and noise was not simulated. (B) From each range of the parameters’ values chosen for the simulation, 7 values were sampled, equally distributed on a logarithmic scale. (C)

Importantly, the average response curve predicted by the model over all parameter combinations that were valid for the males is similar to that predicted for the hermaphrodites with their valid parameter sets (Figure 3F, Figure S3). However, the indication that hermaphrodites and males demonstrate different behaviors suggests that the parameter sets “used” by real worms are more selective. We find that when we simulate the hermaphrodite and male networks using the mutual sets of parameters, the predicted behavior does replicate the distinct behavior of each sex observed experimentally (Figure 3G-H). In other words, we were able to reproduce the dimorphic behavior by using the same parameter sets for the two sexes and changing only the topology, suggesting that the dimorphic topology may by itself account for the behavioral differences.

All possible combinations (823,543) were tested to elucidate whether the activity of the motor neurons in the simulation match the worms’ behaviors in experimental observations. Orange: sets unique to hermaphrodites, cyan: sets unique to males, purple: mutual sets, appropriate for both sexes. (D) Distribution of the biophysical parameters’ values (X axis - seven values per parameter) in the sets that met all the conditions. Color intensity represents the fraction of appropriate sets containing each value (separately for each row). The distribution in the mutual sets is occasionally different, even if the distribution of both sexes is similar. For each parameter, the first row represents all the sets appropriate for the hermaphrodite, the second row is all the sets appropriate for the male, and the third row the mutual sets. Gray represents values that did not appear in the appropriate sets. (E) Principal component analysis was performed on all sets valid for either sex. Data was projected on 2 first principal components (PC1-2). Purple – mutual sets, orange – hermaphrodite-only sets, cyan – male-only sets. (F) Networks using all appropriate sets for each sex do not reproduce the dimorphic response. X axis represents the voltage increase in the sensory neuron following the stimulus (see Methods). Y axis – reverse probability averaged over all sets (18,706 for hermaphrodites, 12,251 for males). Difference in the activation of the motor neurons A and B was taken as an indication for the movement’s direction. (G) Reversal behavior in each of the intersection sets, in response to the sensory stimulus. X axis – voltage increase in the sensory neuron following the stimulus (see Methods). Y axis – number of sets. Purple – both sexes reversed, orange – only the hermaphrodite reversed, cyan – only the male reversed, gray – none of the sexes reversed. (H) Networks using the intersection sets of parameters of both sexes conform with the behavioral differences between them. X axis – voltage increase in the sensory neuron following the stimulus (see Methods). Y axis – reverse probability averaged over all sets (2714 for both sexes).

### Sexually dimorphic interneuron activation in response to a nociceptive cue

Based on the connectivity maps, ASH connections to the interneurons are dimorphic (Figure 2A) and the simulated networks of the two sexes using the parameters of the mutual sets show clear dimorphic activation of the interneurons (Figure 4A). Thus, we used calcium imaging to measure the activity of the AVA interneurons. We found distinct sex-based responses to 0.5 M glycerol in AVA: (1) The peak calcium levels were higher in hermaphrodites than in males (Figure 4B-D). (2) While the calcium signals in hermaphrodites decayed at a slow pace and varied in time, those in the male AVA lasted only while the stimulus was delivered, with levels dropping immediately upon stimulus withdrawal (Figure 4E). As we found the behavioral responses of the two sexes to be similar for strong stimuli (Figure 1C), we also measured the activity of AVA in response to a higher concentration of glycerol. Indeed, at 1 M glycerol, the activity pattern of the male AVA more greatly resembled that of the hermaphrodite (Figure 4F-G) – marked by a comparable peak intensity – but the response duration remained dimorphic (Figure 4H-I). The similar expression levels of the NMDA-type glutamate receptor *nmr-1* in AVA between the sexes (Figure 4J) suggests that the source of the AVA dimorphic activation is not located at the receptor level. Taken together, the predictions of our simulations and the experimental measurements of AVA suggest that the interneuron connectivity plays a key role in the dimorphic behavior.

**Figure 4.**
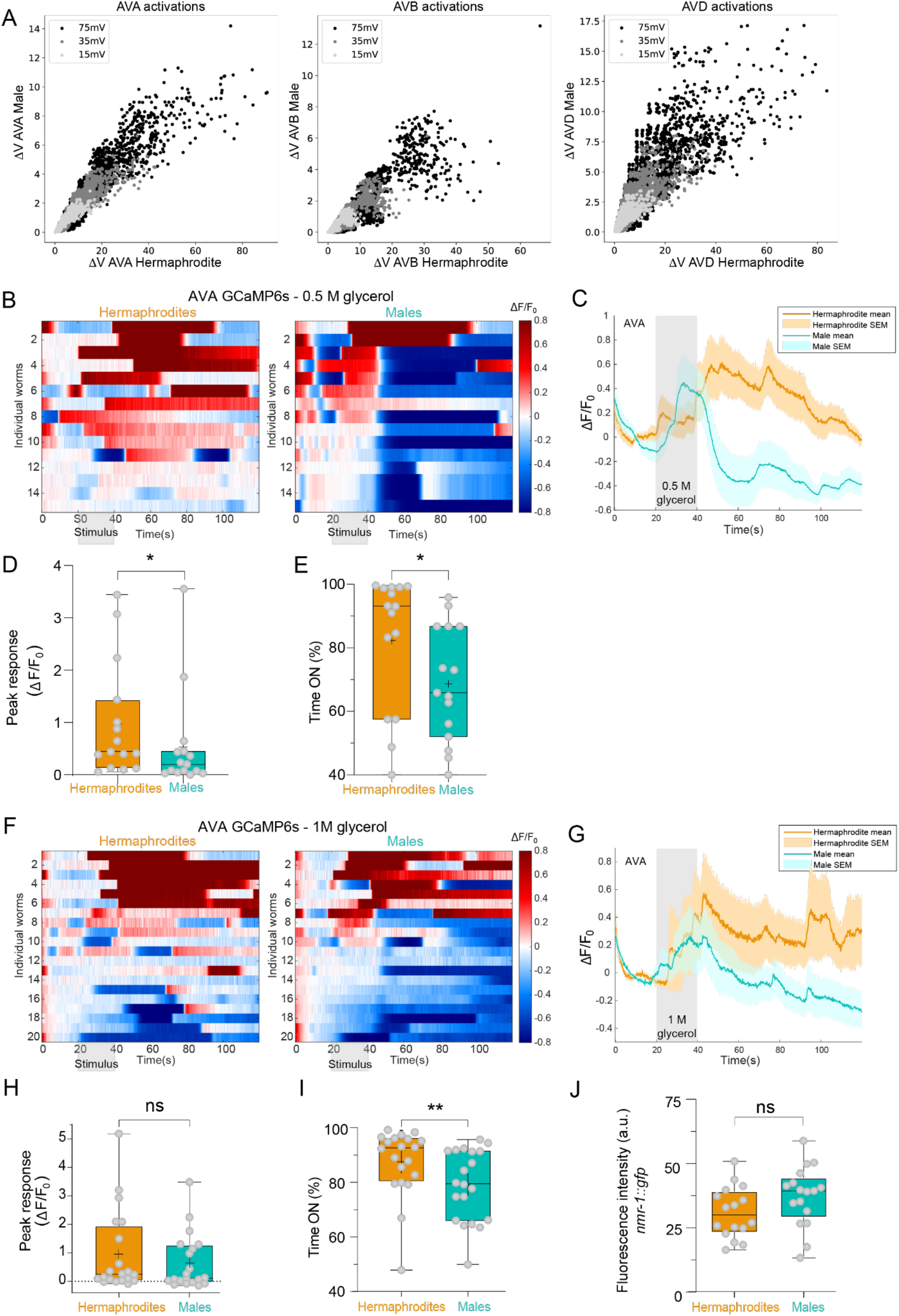
Sexually dimorphic interneuron responses to aversive cues. (A) The change in membrane potential induced by the sensory stimulus in the interneurons directly downstream to ASH in at least one sex (AVA, AVB, AVD, left to right). X axis: The change in the hermaphrodites (mV); Y axis: the change in the males (mV). Each point represents a different parameters’ set out of the mutual sets of both sexes. The strength of the sensory stimulus is represented by the color intensity. (B-I) GCaMP6s calcium responses of AVA interneurons to 0.5 M (B-E, n = 15 animals per sex) and 1 M (F-I, n = 20 animals per sex) glycerol. (B, F) Heatmaps of individual animals representing the normalized calcium levels (color-coded). Stimulus is applied at 20-40 seconds. (C, G) Average and SEM traces of AVA calcium levels. Response intensity (in C) and duration (in C and G) are sexually dimorphic. (D, H) Quantification of peak responses. (E, I) Quantification of response duration. (J) Expression levels of the glutamate receptor *nmr-1* in AVA. n = 14-19 animals per group. D, E, H, I and J are box and whiskers plots, vertical line represents the median and “+” is the mean. We performed a Mann-Whitney test for all comparisons, ** *P*< 0.01, * *P*< 0.05, ns - non-significant.

### Rewiring of single neurons switches behavior in a sexually dimorphic manner

As our data indicated that dimorphic responses in nociception mostly result from differences in connectivity rather than in other neuronal properties, we used our model to identify critical rewiring of the network’s connections that would suffice to switch behavior between the sexes. We simulated “feminization” and “masculinization” of individual neurons in the network (i.e., replacing all the connections of a specific neuron with the connections of the counterpart neuron in the opposite sex, see Methods). Simulating feminization of ASH in males resulted in response curves that resembled the predicted wild-type hermaphrodite response, whereas masculinizing ASH in hermaphrodites had a minor effect on the predicted behavior (Figure 5A middle panel, Figure S4; same results obtained for AVD neuron manipulation, Figure S4B). However, for the AVA neuron, both masculinization and feminization resulted in behavior that resembled that of the opposite sex (Figure 5A right panel; same results received for B motor neuron manipulation, Figure S4E). Thus, our model predicts that changing the sexual identity of individual neurons in the circuit can be sufficient to switch the behavioral response of the worm.

**Figure 5.**
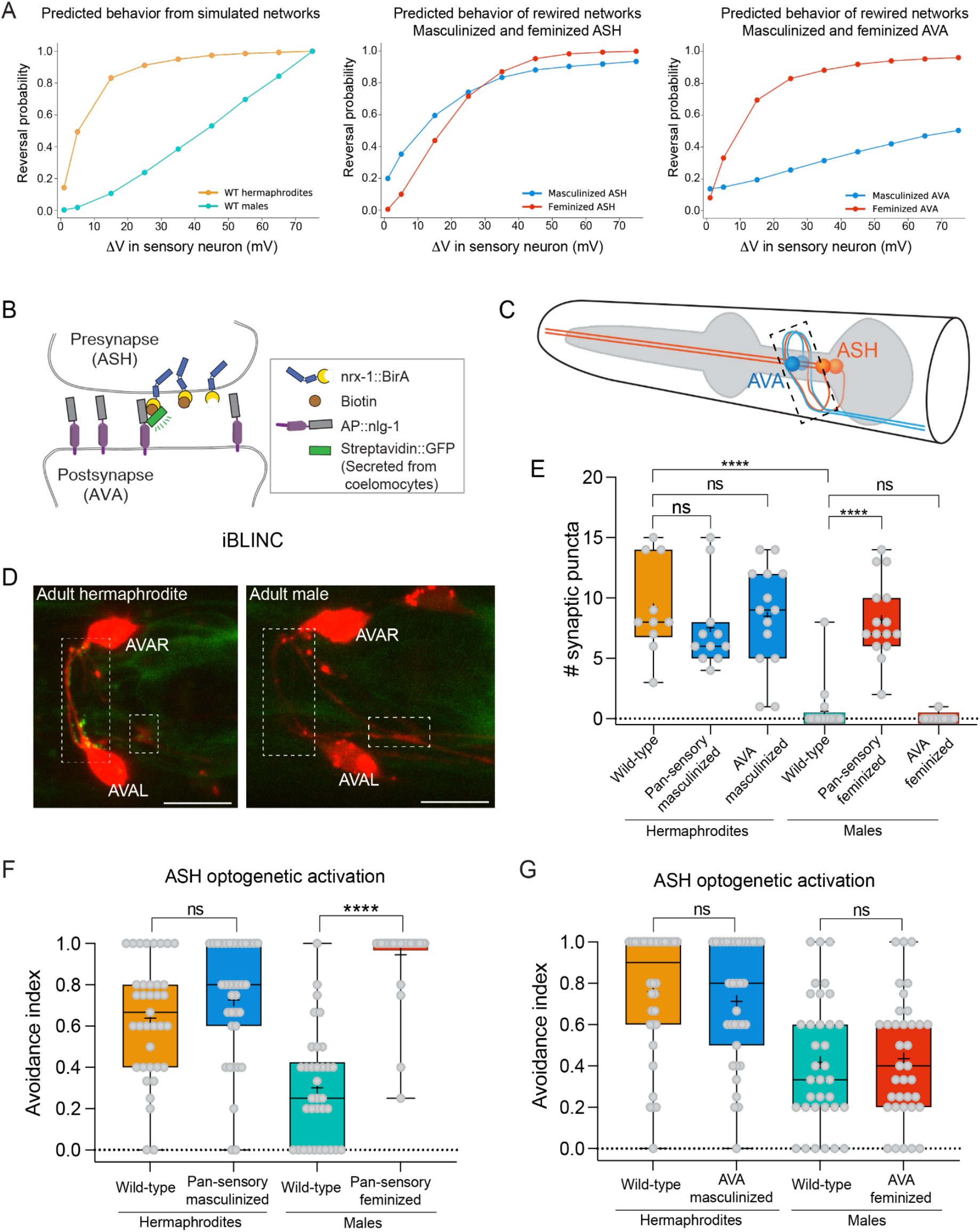
The sexual identity of the sensory neurons shapes the avoidance behavior in a sexually dimorphic manner. (A) Computational model predictions of the escape response in wild-type worms (left), worms with feminized or masculinized ASH (middle), and AVA (right). All connections of ASH and AVA are sex-switched. Backward movement is predicted to increase in ASH-feminized males, yet in ASH-masculinized hermaphrodites, the avoidance is not significantly affected. Backward movement is predicted to increase in AVA-feminized males, and decrease dramatically in AVA-masculinized hermaphrodites. Reverse probability at each voltage point is calculated by averaging the movement direction results of the mutual parameters’ sets of the model. (B) Schematic of the iBLINC technique (*in vivo* Biotin Labeling of INtercellular Contacts) used to visualize and quantify synaptic contacts between neurons (Desbois et al., 2015). (C) Illustration of the contact area (dashed rectangle) between ASH (left and right, orange) and AVA (left and right, blue) neurons at the nerve ring along the ASH axon. (D) Representative confocal micrographs of an adult hermaphrodite and male, showing GFP iBLINC puncta only in hermaphrodites at the contact areas of ASH-AVA (dashed rectangles). AVA neurons are labeled with cytoplasmic mCherry. Scale bar is 10 μm. (E) Quantification of ASH-AVA iBLINC GFP puncta in hermaphrodites and males: wild-type and sex-reversed animals (pan-sensory and AVA, masculinization and feminization, see Methods). n = 10-18 animals per group. We performed a Kruskal-Wallis test followed by Dunn’s multiple comparison test, **** *P*< 0.0001, ns - non-significant. (F) Avoidance index for ASH optogenetic activation in wild-type or pan-sensory sex-reversed animals. In males, but not in hermaphrodites, changing the sex of the ciliated sensory neurons is sufficient to flip the avoidance behavior, in agreement with the model’s predictions. LED intensity is ~1.47 mW/mm^2^. n = 33-37 animals per group. (G) ASH optogenetic activation in wild-type worms and worms with masculinized or feminized AVA. LED intensity is ~1.47 mW/mm2. n = 34-36 animals per group. E, F and G are box and whiskers plots, vertical line represents the median and “+” is the mean. We performed a Mann-Whitney test for all comparisons, **** *P*< 0.0001, ns - non-significant.

We then turned to test these predictions experimentally by manipulating the sex-determination pathway of the worm to “feminize” (*tra-2* expression) or “masculinize” (*fem-3* expression) specific neurons (Mehra et al., 1999; Mowrey et al., 2014; Oren-Suissa et al., 2016)(See Methods). To verify that the sex-reversal manipulations resulted in the desired synaptic connectivity changes, we trans-synaptically labeled the connection between ASH and AVA using *in-vivo* Biotin Labeling of INtracellular Contact (iBLINC, Figure 5B-D) (Desbois et al., 2015). First, we analyzed GFP puncta of the ASH-AVA connection temporally in wild-type animals, and confirmed that the connection is hermaphrodite-specific at the adult stage (Figure S5A). Second, we analyzed the connection in sex-reversed animals, and found, surprisingly, that in hermaphrodites, sex-reversal experiments did not result in changes to ASH-AVA connectivity. However, in males, pan-sensory feminization resulted in the formation of ectopic ASH-AVA synapses (Figure 5E).

We then tested whether the sex-reversal of single neurons changed the behavior induced by optogenetic stimulation of animals expressing ChR2 in the sensory ASH neuron. We found that, indeed, pan-sensory feminization resulted in sex-reversal of the male behavior (Figure 5F), in agreement with our model’s prediction. As our synaptic measurement showed that masculinization of ASH did not change the desired synaptic connection in the hermaphrodite, or in sex-reversal of AVA, it was reassuring to find that these attempted manipulations did not change the worms’ behavior (Figure 5F,G).

Our transsynaptic labeling between ASH and AVA enabled us to assess the developmental aspect of these dimorphic connections. Synapses are first observed at the 3rd larval stage in both sexes, but then synapses are sex-specifically removed in males (Figure S5A). In line with the non-dimorphic synaptic connectivity at this stage, juvenile animals (L3) respond in a non-dimorphic manner to the nociceptive 0.5 M glycerol stimulus (Figure S5B), suggesting that the sexually dimorphic response to aversive stimuli is unique to the adult nervous system. Taken together, our results suggest that the hermaphrodites’ connectivity map and resulting behavior are more robust compared to those of males.

### Rewiring a single synaptic connection is sufficient to flip behavior to that of the opposite sex

Given the success of changing behavior by sex reversal at the neuronal level, we turned to ask what would be the behavioral effect of rewiring individual synapses. To identify which specific synaptic connections are crucial for generating the dimorphic nociceptive behavior, we used our simulated networks to explore the potential effect of flipping single synaptic connections or pairs of connections simultaneously. Our simulations predict that removing hermaphrodite-specific connections would not induce a male-like response, but may have a small effect in the case of removing pairs of connections (For example, ASH-AVA)(Figure 6A and S6). Conversely, the addition of hermaphrodite-specific connections to males can induce a behavioral switch (Figure 6B, Figure S6). Neither sex is predicted to be affected by the removal or addition of male-specific connections (Figure 6S). Following these predictions, we manipulated the ASH-AVA connection experimentally. We generated a transgenic strain in which we inserted a mammalian connexin36-mediated synthetic gap junction (Rabinowitch et al., 2014) between both neurons (Figure 6C). ASH-connexin36 puncta coincided with AVA-connexin36 puncta, and localized along the overlapping region between the two neurons at the nerve ring (Figure 6D), as expected. We assayed the behavior of these males in comparison to wild-type males and found that the specific rewiring increased the probability of reversal to a level similar to that of wild-type hermaphrodites, in agreement with our model (Figure 6E, Figure 1C).

**Figure 6.**
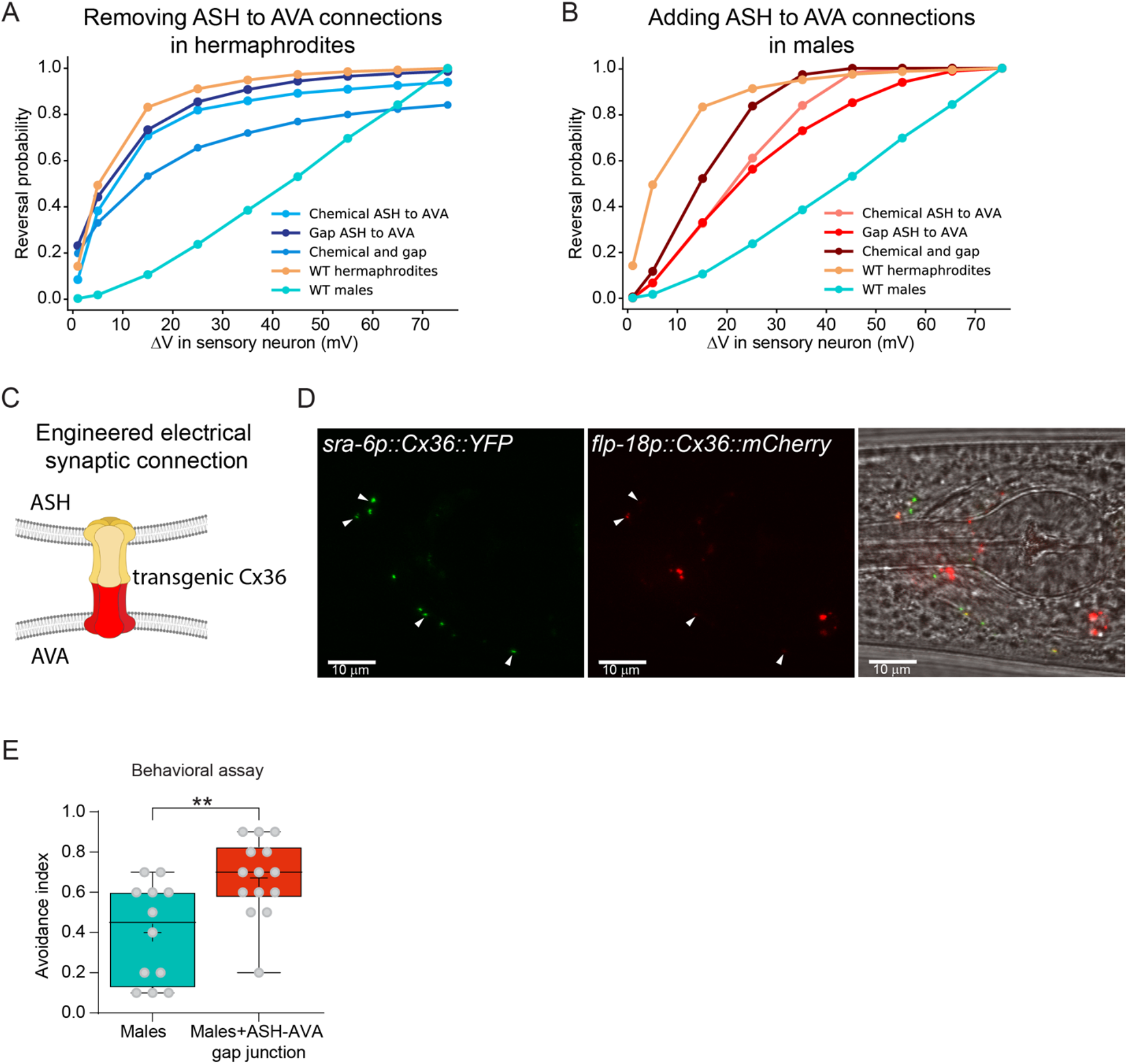
Rewiring of a single connection in the circuit is sufficient to feminize male avoidance behavior. (A) Model predictions of the escape response in wild-type worms, and hermaphrodites without the ASH-AVA connection (chemical, electrical, or both). Backward movement is predicted to slightly decrease in hermaphrodites without ASH-AVA connections. (B) Model predictions of the escape response in wild-type worms, and in males with an additional connection between ASH and AVA (chemical, electrical, or both). Backward movement is predicted to increase in males with ASH-AVA connections. The reversal probability at each voltage point is calculated by averaging the movement results of the mutual parameters’ sets of the model. (C) Illustration of an engineered electrical synapse. A fluorescently tagged codon-optimized mouse connexin Cx36 is expressed under the ASH and AVA promoters (tagged with YFP and mCherry, respectively) (Rabinowitch et al., 2014). (D) Representative confocal micrographs of a male expressing a synthetic ASH-AVA electrical synapse. *ASHp::Cx36::YFP* and *AVAp::Cx36::mCherry* puncta (white arrowheads) colocalize along the contact area of ASH andAVA. Scale bar is 10 μm. (E) In a tail drop assay with a 0.5 M glycerol stimulus, the backward movement of males with a gap junction between ASH and AVA is significantly stronger than that of wild-type males. E is a box and whiskers plot, vertical line represents the median and “+” is the mean. n = 12 wild-type males, 14 males with ASH-AVA synapse. We performed a Mann-Whitney test, ** *P*< 0.01.

### Rewired males are less successful in finding a mate

Our rewiring experiments enabled us to explore an even wider behavioral implication of the topology of the nociceptive network: We asked how the circuit’s rewiring might affect male behavior in a way that could be translated into environmental conditions met in nature. To address this question, we created a setup in which a single male is placed in a compound environment containing a confined group of hermaphrodites, which served as the positive sensory cues, while receiving repeated nociceptive stimuli, delivered optogenetically (Figure 7A). We compared the behavior of wild-type males expressing ChR2 in ASH (in which ASH activation generates a significantly reduced escape response compared to hermaphrodites, see Figure 2D) to that of similar males but with feminized sensory neurons (i.e., that respond in a hermaphrodite-like manner to the optogenetic stimulation, Figure 5F). We found that activating ASH in sensory-feminized males significantly extended the time required for them to reach the hermaphrodites (Figure 7B-C). Importantly, the control group of pan-sensory feminized males without the activation co-factor ATR showed normal attraction towards the hermaphrodites. These results suggest that having a topology that enables efficient avoidance of noxious cues in males would incur a “cost” in terms of finding mates.

**Figure 7.**
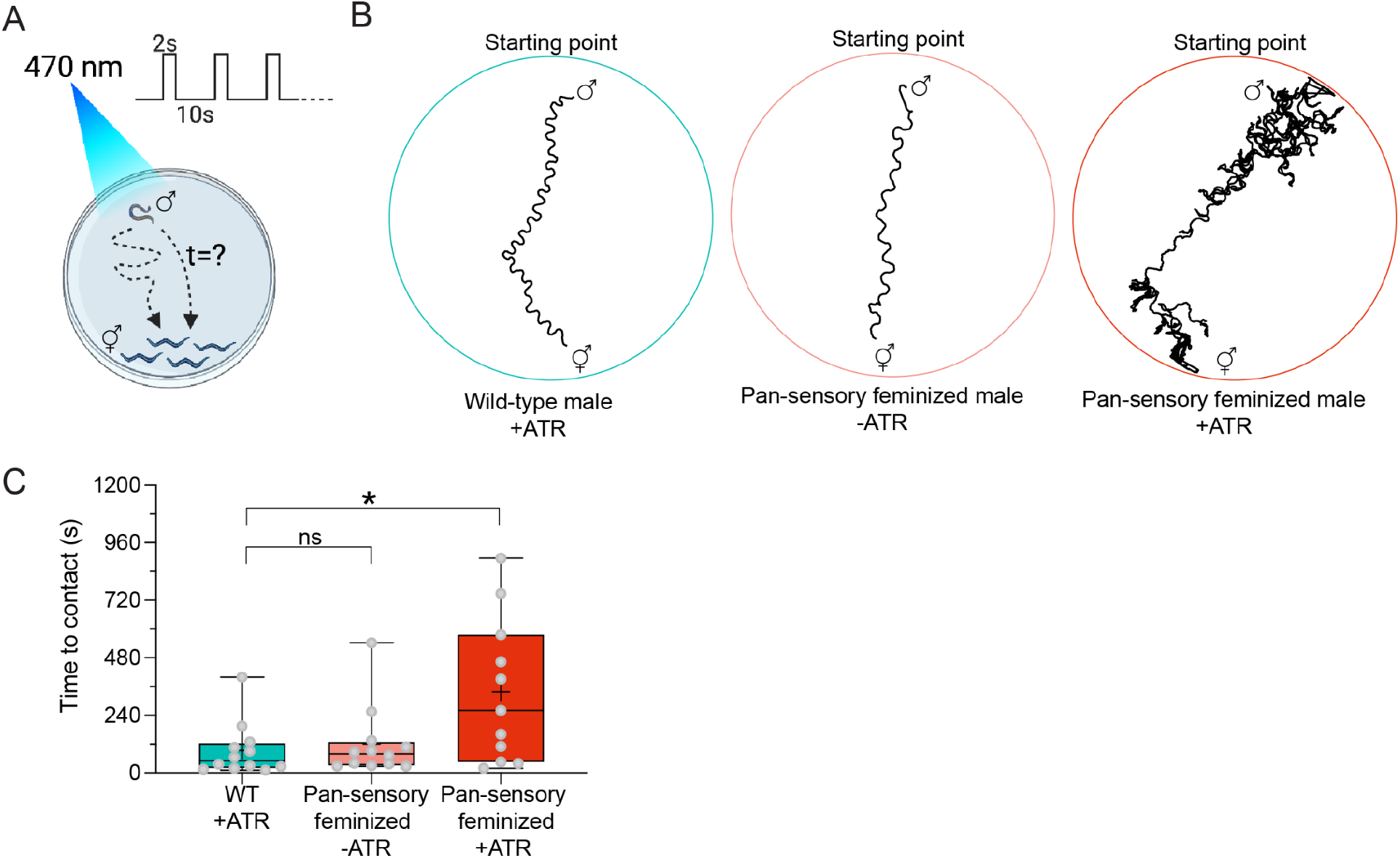
Feminized avoidance behavior impedes males’ ability to reach a mate. (A) Illustration of the experimental design. Repetitive optogenetic stimulations of ASH are applied while the male is searching for the immobilized hermaphrodites. The time it takes the male to contact a hermaphrodite is quantified (see Methods). (B) Representative traces of a WT male +ATR (left), a pan-sensory feminized male -ATR (middle), and a pan-sensory feminized male +ATR (right). (C) Quantification of the time to contact of WT males with ATR, pan-sensory feminized males without ATR, and pan-sensory feminized males with ATR. n = 11-12 males per group. C is a box and whiskers plot, vertical line represents the median and “+” is the mean. We performed a Kruskal-Wallis test followed by Dunn’s multiple comparison test, * *P*< 0.05, ns - nonsignificant.

## DISCUSSION

We explored the design of a sex-shared neural circuit that controls the fundamental trait of response to pain sensation. We found that *C. elegans* exhibits sexually-dimorphic nociceptive behaviors, and that these behaviors are mediated by a compact neuronal circuit of sex-shared neurons that are connected in a dimorphic manner. We further demonstrated that differences in circuit connectivity, rather than in sensory transduction, shape the sex-specific behavior. We extensively analyzed the sensory response to the shared stimuli. Such direct measurements of the processing at the sensory level itself have been often neglected in studies deciphering sexually dimorphic behaviors (Fan et al., 2018; Keddy-Hector et al., 1992). Instead, much attention has been directed to differences downstream to the sensory level (Chiu et al., 2021; Hoke et al., 2010; Sten et al., 2021). Even though some sex-specific olfactory ligands have been identified, the neural mechanisms that enable a sexually-dimorphic response to these odors remain largely unknown (Stowers and Logan, 2010).

Our simulations of the nociceptive circuits identified a relatively narrow range of parameters that were valid for both male and hermaphrodite circuits. Importantly, these sets of parameters replicated the experimentally observed dimorphic behavior, despite the simplifications made by the model (i.e., the model did not consider noise, inputs from sex-specific neurons or other circuits, all the neurons were simulated as excitatory, and possible variability between cells within each animal was neglected). While it is possible that males and hermaphrodites may still use different values, these sets suggest sex-agnostic biophysical parameters for this circuit (i.e., network parameters do not require sex-dependent regulation (see (Marder and Taylor, 2011)) and that the dimorphic topology alone can account for the behavioral differences.

Our model allowed us to identify which dimorphic connectivity structures shape behavior, and how to change connectivity to reprogram behavior. These predictions were then validated experimentally, and revealed that the hermaphrodite circuit is more robust to changes than the male one. This surprising result is in contrast to previous studies that used reciprocal sex-reversals (i.e., both female-to-male and male-to-female) to demonstrate that neuronal sexual identity functions cell-autonomously (Fagan et al., 2018; Mowrey et al., 2014; Oren-Suissa et al., 2016; Serrano-Saiz et al., 2017; Wu et al., 2018). Our non-reciprocal results may point to non-autonomous effects that are mediated either by suppressive signals coming from the hermaphrodite body or the absence of positive signals coming from the male body. Alternatively, a cell-autonomous signal independent of the known sex-determination pathway could control the hermaphrodite connectivity. The hermaphrodite circuit’s greater robustness to changes and the presence of mechanisms that maintain this robustness suggest that the hermaphrodites’ tendency to respond more to aversive stimuli is an important property.

Calcium imaging of the responses to aversive stimuli of the AVA interneuron revealed variability in the activity of this neuron, and some males exhibited hermaphrodite-like behavior (Figure 4B). Such inter-individual and sex-dependent differences have been reported for different organisms (Goaillard and Marder, 2021; Kfir et al., 2012; Körholz et al., 2018; Schneidman et al., 2000; Zajitschek et al., 2020), but the underlying molecular and cellular mechanisms generating behavioral individuality remain unclear. The hermaphrodite-like activity of this neuron in some of the males may arise from variability in connectivity between individuals. This suggests that the behavioral differences among individuals might have a topological source (e.g., a weak connection between ASH-AVA might persist in these unique males), and could also reflect developmental and evolutionary implications.

Sexual selection describes a set of dynamics operating within species that are the evolutionary drivers of sex differences. Sexual selection and natural selection are viewed as distinct evolutionary forces, occasionally even acting in a contradictory manner in cases where sexual selection selects for phenotypes that are not favored by natural selection (Hosken and House, 2011; Mead and Arnold, 2004). We speculate that there is a sexually-dimorphic need to avoid aversive stimuli; this notion is supported by our behavioral experiments in female-male *Caenorhabditis* species. In males, sexual selection acts against the evolutionary pressure to avoid hazardous substances by modifying the circuit in favor of the mating circuit, resulting in connectivity differences in the sex-shared nociceptive circuit. Supporting this hypothesis, reprogrammed males demonstrate increased sensitivity to aversive stimuli, as well as decreased efficiency in making contact with hermaphrodites, suggesting that the male would suffer a physiological “cost” for a topology that enables efficient avoidance of noxious cues. Our results demonstrate that subtle topological changes in neuronal circuits are sufficient to change their function, which is likely to play a key role in the adaptation to different environmental and sexual needs.

## ACKNOWLEDGEMENTS

We thank members of the Oren-Suissa lab and Schneidman lab for their critical insights regarding the manuscript, Eduard Bokman for help and discussions on calcium imaging, Cori Bargmann, William Schafer, Shai Shaham, Menachem Katz, Ithai Rabinowitch and Oliver Hobert for *C. elegans* strains and constructs. Some strains were provided by the CGC, which is funded by the NIH Office of Research Infrastructure Programs (P40 OD010440).

This work was supported by the European Research Council ERC-2019-STG 850784 (MOS), Israel Science Foundation grant 961/21 (MOS), the Simons Collaboration on the Global Brain Grant 542997 (ES), Martin Kushner Schnur and Mr. and Mrs. Lawrence Feis (ES). This research was also partially supported by the Israeli Council for Higher Education (CHE) via the Weizmann Data Science Research Center (MOS and ES). MOS is grateful to the Azrieli Foundation for the award of an Azrieli Fellowship, and is the incumbent of the Jenna and Julia Birnbach Family Career Development Chair. ES is the incumbent of the Joseph and Bessie Feinberg Chair.

## AUTHOR CONTRIBUTIONS

VP, YS conducted the experiments, AC, RAB, JRH, MLE, RMM, NJR contributed to behavioral experiments (“Head drop assays”). GG carried out all simulations and modeling experiments. DSP, DMF, ES and MOS supervised and designed experiments. VP, YS, GG, ES and MOS wrote the paper.

## METHODS

### Repellent assay - tail drop

Tail drop avoidance assay was described previously (Hilliard et al., 2002; Oren-Suissa et al., 2016). All assays were done on 1day adults. Briefly, worms were given ten minutes to habituate on a foodless NGM experiment plate, and then underwent 8-10 repellent stimulations with at least 2 minute intervals between stimuli. A small drop of the repellent (glycerol in S basal, SDS or copper (CuSO4) in M13 buffer (30 mM Tris-HCl (pH 7.0), 100 mM NaCl, 10 mM KCl) was placed on the agar near the tail of a forward-moving animal, using a 10 μl glass-calibrated pipette (VWR International), pulled by hand on a flame to create two needles with reduced diameter. The pipette was mounted in a holder with a rubber tube, operated by mouth. A day before the experiment, unseeded NGM plates were taken out of storage at 4°C, dried for 2 h at 37°C, and then left on the bench. Scoring for each trial was binary (1 for reversal, 0 for no reversal). The average score of all trials is the avoidance index for each animal. For the tail drop assay of L3 animals, worms were kept separately following the assay until their sex was identifiable.

### Repellent assay - head drop

The head drop assay was performed for the repellents glycerol, SDS, copper and quinine, in M13 buffer following the protocol described by Hilliard et al. (Hilliard et al., 2002, 2004). In each experiment, ~15 young adult worms were placed on a dry unseeded NGM plate and given ten minutes to habituate before each worm was subjected to one head-drop test. Namely, a small drop of stimulus was placed in front of a forward-moving worm, which was then scored according to its movement direction during a 4 sec window. The drop was placed with a fresh capillary pulled on a flame. Each assay plate received a score according to the calculation: (Number of worms reversed)*100/(Total number of worms tested on the plate). For each experiment, assays were repeated on at least 3 independent days.

### Microfluidic chip fabrication and calcium imaging

Olfactory chips were fabricated according to (Chronis et al., 2007) with the help of the Nanofabrication Unit at the Weizmann Institute of Science. In short, a polydimethylsiloxane (PDMS) mixture was cast into premade 0.5-cm-high chip molds and allowed to solidify at 65°C for 3 h. Individual chips were cut by hand with a scalpel and then punctured to create fluidic inlets using a PDMS biopsy punch (Elveflow) 0.5 mm in diameter. The chips were attached to glass coverslips by exposing them to plasma for 30 s, then manually attaching them together and drying them on a hot plate for 1 h at 65°C. The tunnel height was 28 μM and the width at the worm’s nose space was 24 μM.

The microfluidic chip was operated using two pumps that control the flow of buffer and stimulus into the microfluidic chip. The solutions were pushed through PVC tubes and stainless-steel connectors into the tunnels of the chip, and with a manual switch, we determined the arrival of the stimulus to the worm. Tubes were replaced between experiments and connectors cleaned with ethanol. The pumping rate during experiments was ~0.005 ml/min. Loading the worm into the chip was done by placing the worm in a drop of S basal buffer, sucking the drop with a 1 ml syringe and inserting it into the relevant inlet of the chip. After proper worm positioning, 2 minutes for habituation were given prior to imaging with the lasers on. To prevent movement, 10 mM Levamisole was added to all solutions (except for the S basal solution used to load the worm). To visualize proper delivery of the stimulus to the worm, 50 μM rhodamine B was added only to the stimulus. If the worm moved or the flow was incorrect, the file was discarded and a second trial was performed with the same worm. No more than two trials were done with the same worm. Imaging was done with a Zeiss LSM 880 confocal microscope using a 40x magnification water objective. Imaging rate was 6.667 Hz, total imaging duration was 2 min, and stimulus duration was 20 sec. Stimulus was given at 20 to 40 sec from imaging initiation. For analysis, the GCaMP6s fluorescence intensity was measured using FIJI. All files were exported as tiff files, ROIs (regions of interest) of the somas were drawn manually to best represent the signal, and their mean gray values were exported. Downstream data processing was performed using MATLAB. For each worm, the baseline fluorescent level (F_0_) was calculated by averaging the mean gray values of 100 frames (15 sec) before stimulus delivery. Then, for each frame, the ΔF was calculated by subtracting F_0_ from the value of that time point, and the result was divided by F_0_, to normalize differences in fluorescence baseline levels between individuals (ΔF/F_0_).

All statistical comparisons were done on the normalized data. For peak response comparisons, the maximal values of each worm from 20 to 60 sec of imaging were used. Response duration comparisons were done as in (Pradhan et al., 2019). In short, first, the response range was calculated for each worm (subtraction of the minimum value from the maximum value obtained in the time window of 20 to 60 sec of imaging). Then, the number of frames with above-threshold values were quantified for each worm (if value_i_ ≥ minimum value_i_ + threshold * range_i_), using a 5% threshold, and converted into percentages.

### pHluorin imaging

One-day adult worms expressing sra-6::eat-4::pHluorin (Ventimiglia and Bargmann, 2017) were imaged in a microfluidic chip and a 40X magnification water objective of a confocal microscope, similarly to the calcium imaging procedure. Each animal was imaged with a z-stack of 28 slices, which was ~14.78 μm thick. The duration of imaging was 210 sec or 35 frames, at an imaging rate of 0.1666 Hz, and the 0.5 M glycerol stimuli were given at 36-60 sec / 6-10 frame, 96-120 sec / 16-20 frame and 156-180 sec / 26-30 frame. The analysis was partially automated; all images were first processed to delete z-slices that did not contain any relevant signal, and then, all the remaining z-slices were summed and cropped in FIJI to retain only the axon of ASHL/ASHR. To resolve any movements of the worms, the movies were registered with the StackReg plugin using a rigid body transformation. The ROIs were then drawn manually, and mean gray values were extracted. The ΔF/F_0_ normalization was performed with MATLAB, and the baseline fluorescent level (F_0_) was calculated for each stimulus separately using the 4 frames before each stimulus.

### Optogenetics

We used worms that only express ChR2 in ASH neurons, using the intersectional approach^27^ (*gpa-13p::FLPase, sra-6p::FTF::ChR2::YFP*), and mutated in the *lite-1* gene, to ensure the measured behavioral responses are not due to the activity of the endogenous blue-light receptors, but due to ChR2 activity. Worms at the L4 developmental stage were selected a day before the experiment and separated into hermaphrodite and male control and experiment groups. They were transferred to newly seeded plates with 300 μl OP50 that was concentrated at 1:10. ATR was added only to the experiment groups’ plates, to a final concentration of 100 μM ATR. As ATR is sensitive to light, all the plates were handled in the dark. Tracking and optogenetics were done on unseeded NGM plates. The day before the experiment, the plates were taken out of storage at 4°C, dried for 2 h in 37°C, and left on the bench overnight. On the experiment day, these plates were seeded with 30 μl OP50 (and ATR, only for the experiment group’s plates). Experiments were performed at 24°C. To keep the worms in the camera’s field of view, a plastic ring was placed on the agar and five worms were placed inside it. After 10 minutes of habituation, the worms were tracked for 69 sec with five 2-sec LED activations each, and 10 sec ISI. Recording started with 10 sec without LED. Analysis was performed manually. If the worm reversed during a 3 sec window (2 sec LED duration + one additional second), it received a score of one, otherwise a score of zero. The five results of each worm were averaged to a number between zero and one. If the worm touched the ring or was moving backwards while the LED turned on, the trial was not counted.

### DiO/DiD staining

Worms were washed with M9 buffer and incubated for 1 h in 1 ml M9 and 5 μl DiO/DiD dye in ~25 rpm tilt. The worms were then transferred to a fresh plate and let to crawl on a bacterial lawn.

### Imaging

Prior to imaging, worms were mounted on a 5 % agarose pad on a glass slide, on a drop of M9 containing 100 mM sodium azide, which was used as an anesthetic. A Zeiss LSM 880 confocal microscope was used with 63x magnification, and all imaging parameters were kept identical in each imaging experiment. For the EAT-4/VGLUT expression measurement, ASH was identified based on its morphological position relative to all the stained DiO neurons, and the z-plane with the strongest signal was exported to measure fluorescence intensity using FIJI. ROIs were manually drawn.

### Attraction assay and optogenetics

All preparations for the attraction assay were identical to the preparations of the optogenetic experiments. 2 μl of 10 mM sodium azide were placed on a dried, unseeded NGM plate and allowed to dry for ~5 minutes. Then, a plastic ring ~5 mm in diameter was placed on the plate, and 15 *unc-31* hermaphrodites were placed in the area with the absorbed sodium azide. A single male was placed each time inside the ring on the side opposite to the location of the hermaphrodites, and immediately a recording started with blue LED activations (10 seconds ISI, 2 second duration). When the male touched a hermaphrodite, the recording and optogenetic activations were terminated, the male was removed from the plate, and a new male was added for the next trial. The time it took the male to reach the hermaphrodite served as its score for the statistical comparison. If the male did not reach a hermaphrodite within 20 minutes, the trial was terminated and discarded from the statistics.

### Molecular cloning

The intersectional approach was employed to achieve ASH-specific expression (Ezcurra et al., 2011). pTNZ141 (*gpa-13p::FLPase*) was injected with a version of pTNZ109 (*sra-6p::FTF::ChR2::YFP*) in which the ChR2::YFP cassette was replaced with the transgene of interest. For calcium imaging of ASH, ChR2::YFP was replaced with GCaMP6s, amplified from p45.641 (*mec-4p-nls-RSET-GCaMP3(CEopt)-SL2-nls-TagRFP-unc-54utr*, a kind gift from Doug Kim).

Pan-sensory masculinization and feminization was achieved in a set of ciliated neurons by expressing *tra-2[intracellular]* (Fagan et al., 2018) or *fem-3* under the *osm-5p*.

For calcium imaging of AVA, the GCaMP6s fragment was amplified from p45.641 and inserted into a backbone containing the *flp-18p* promoter by Gibson assembly. See STAR methods for primers.

To create the iBlinc synaptic labeling between ASH and AVA, a *sra-6p* promoter fragment (for the ASH side) was linked to the birA::nrx-1 backbone by Gibson assembly. The resulting plasmid was injected with the plasmid pMO26 (*flp-18p::AP::NLG-1*) for the AVA side.

To generate an artificial gap junction between ASH and AVA, a *flp-18p* promoter fragment (for the AVA side) was inserted instead of the *gpa-11p* promoter in plasmid pIR204 (*gpa-11p::Cx36::mCherry*) by Gibson assembly. The resulting plasmid was injected with the plasmid pIR111 encoding for ASH-specific Cx36, *sra-6p::Cx36::YFP* (pIR111 and pIR204 were a kind gift from Ithai Rabinowitch).

### Extracting connectivity data

Connectivity data was taken from (Cook et al., 2019). The strength of connectivity between each pair of neurons was described as the total number of electron-microscopy sections containing that connection. For the simulation, we used the average of the corresponding synaptic connections of the neurons in the left side and right side of the worm from (Cook et al., 2019), and each neuronal class (A and B neurons) were treated as a single cell (Rakowski et al., 2013). Connections within a neuronal class or between corresponding neurons on the left and right circuits were ignored. Chemical synapses between neural groups or pairs were simulated if it appeared in more than 10 serial sections. We note that these criteria neglect possible asymmetry between the right and left side of the worm, as well as differences between the ventral and dorsal side, and that lumping together all neurons of the same class, does neglect possible structure within the class.

### Simulating membrane potential of neurons in the nociceptive circuit

The membrane potential of the neurons in the circuit was calculated by (Gerstner et al., 2009; Varshney et al., 2011):

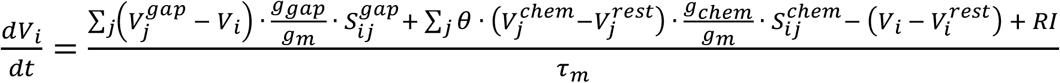
where *V_i_* is the voltage of the postsynaptic neuron, and *V_j_* is that of its presynaptic partners, and the sums are over all neurons in the circuit. *rest* denotes the membrane potential at the resting state, *chem* and *gap* denote chemical synapses and gap junctions, respectively. *g* denotes conductance, and *g_m_* is the membrane conductance of neuron *i*. *S_ij_* is the strength of connection between neuron *i* and neuron *j*. *R* is the resistance of the neuron, *I* is the external current, and *τ_m_* is the membrane time constant. *θ* is an indicator of the activity of chemical synapses:

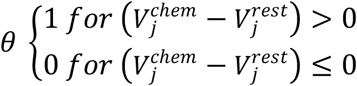
such that chemical synapses are active only when the presynaptic cell is activated, (i.e., above its resting potential). The neurons in this circuit are assumed to release neurotransmitters tonically and to not use action potentials (Varshney et al., 2011). We, therefore, simulated the effect of a chemical synapse as proportional to the strength of activation of the presynaptic cell (voltage above its rest value). We simulated the membrane potential of the neurons for 15 seconds, with time steps Δt = 0.2 ms. The sensory stimulus was presented as an external input current to the head sensory neuron ASH for 5 sec, starting from the 5th second of the simulation. To replicate the spontaneous forward movement of the worms, we assumed a basal activation of interneurons AVA, AVB, PVC – as these neurons’ spontaneous activity correlated with the direction of the movement (Kawano et al., 2011; Li et al., 2011; Wicks and Rankin, 1997). The basal inputs to the interneurons were “on” throughout the simulation.

As the values of different biophysical parameters of the neurons in this circuit are unknown, we used 7 different parameters for the simulations. For each parameter, we used 7 different values that were equally spaced on a logarithmic scale from a biologically reasonable range (Goodman et al., 1998; Lindsay et al., 2011; Palyanov et al., 2012; Rakowski and Karbowski, 2017; Rakowski et al., 2013; Roehrig, 1998; Varshney et al., 2011; Wicks et al., 1996). Rm values were within 15kΩ-1500kΩ, capacitance values were 0.1uF-10uF, values of conductance of gap junctions and chemical synapses were 1pS-1nS, and the basal input values of the interneurons triggered a voltage change of 1mV-75mV. Overall, we explored 7^7^=823,543 parameter combinations. The specific parameters of all the neurons in the circuit were assumed to be the same. *R_in_* was calculated separately for each set of parameters, using the changing value of *R_m_* and a fixed value of the surface area of a cell, which was taken to be 15 · *10^−6^cm^2^* (Goodman et al., 1998; Palyanov et al., 2012; Rakowski et al., 2013; Varshney et al., 2011; Wicks et al., 1996). To induce a voltage-change of a desired magnitude, we calculated the currents to the sensory neurons and to the interneurons, using the values of each parameter set.

### Manipulation of synaptic strength values

For each parameter set, the synaptic strength values were randomly and independently strengthened in 20%, weakened in 20%, with the remainder left unchanged.

### Assessing the validity of each parameter set

All combinations of the parameter values were evaluated to determine whether they would result in behavior that matched that of the experimentally observed one. To qualify, the parameter set had to meet the following conditions: First, the membrane potential of all the neurons had to remain lower than 100mV throughout the simulation, and the time constant had to be lower than 500 ms. Second, the membrane potential of all the neurons had to move back toward the resting potential once the stimulus was over (namely, the difference between the membrane potential of all the neurons and their resting potentials 5 sec after the stimulus was over had to be smaller than that at the termination time of the sensory stimulus). Since in the absence of external stimuli worms typically move forward, we required the membrane potential of motor neuron B in the absence of a sensory stimulus to be higher than that of motor neuron A, which would result in a forward movement. Accordingly, we required backward movement in our simulation (A higher than B), after a strong sensory input. While we simulated the response of the circuit to 9 strengths of sensory stimuli to ASH, the requirements above were applied only to the strongest one.

### Assessing the simulated response of the interneurons

For the sets of parameters that were valid for both the males and hermaphrodites, we focused on stimuli that induced specific changes to the membrane potential of the sensory neurons (75mV, 35mV, 15mV). For each set and stimulus value, we measured the resulting change in membrane potential in the interneurons AVA, AVB, AVD. The change was measured as the difference between the maximal membrane potential value within the time of the stimulus and the membrane potential one time point before the stimulus began.

### Perturbations to the circuit’s connectivity

For each of the sets that were valid for both the males and the hermaphrodites, we simulated three kinds of perturbations: removing or adding a single dimorphic connection, removing or adding two dimorphic connections simultaneously, or replacing the connectivity of an individual neuron (here all connections of a specific cell were copied from the opposite sex - incoming and outgoing synapses and gap junctions); non-dimorphic connections were altered as well, and received the strength of the connection in the opposite sex. For each perturbation, we examined the response of the simulated circuits to multiple strengths of sensory stimuli.

