## Supplemental information for "Sex-specific topology of the nociceptive circuit shapes dimorphic behavior in *C. elegans*"

### SUPPLEMENTAL FIGURES

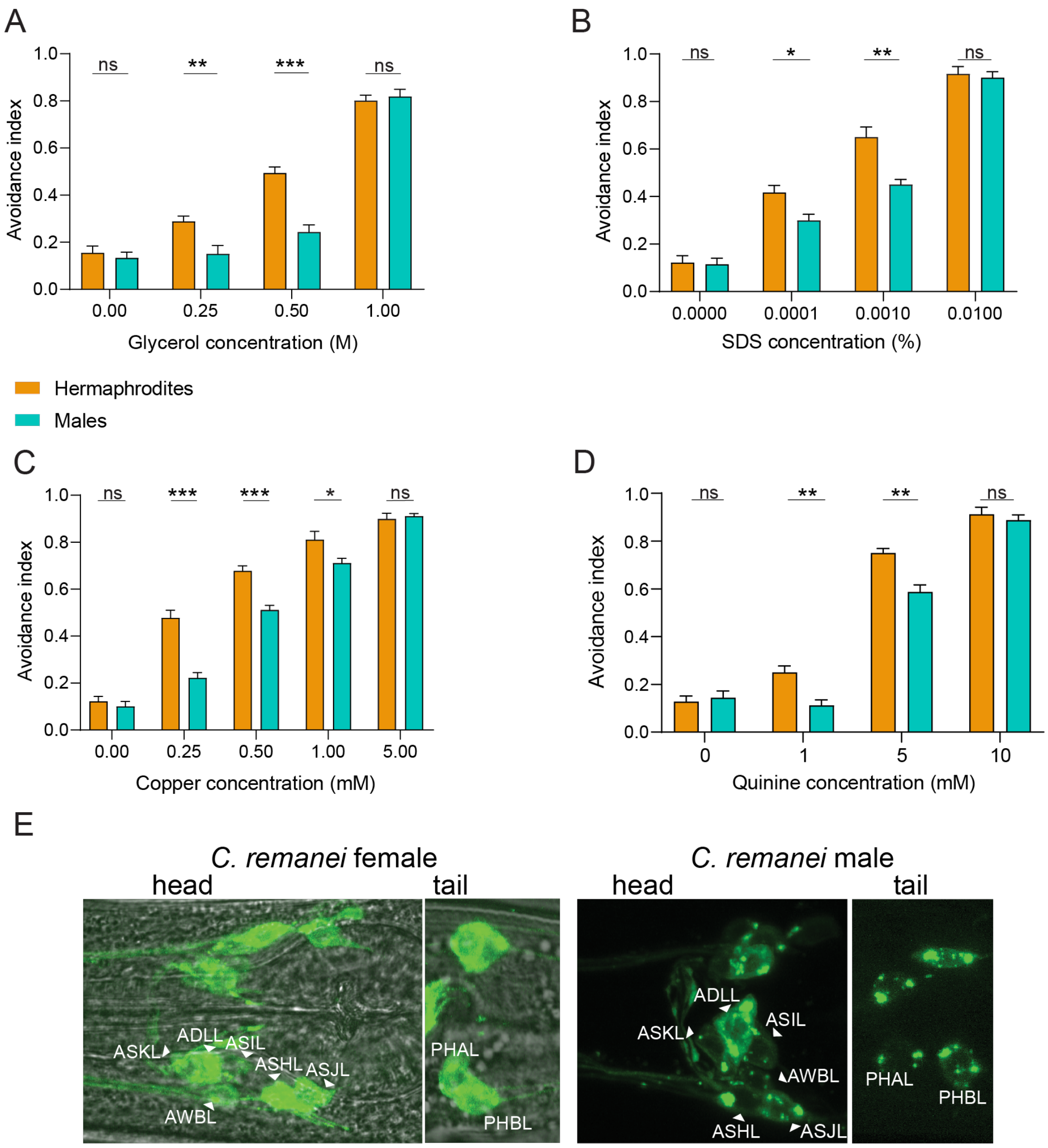


**Figure S1. Sexually dimorphic avoidance behavior to multimodal stimuli**

(A-D) Chemosensory repulsion head-drop assays (see Methods for full description). Hermaphrodites reverse more than males at intermediate concentrations of aversive stimuli - glycerol (A), SDS (B), copper (C) and quinine (D). n = glycerol - 9 repeats per concentration, SDS - 6-9 repeats per concentration, copper - 9 repeats per concentration, quinine - 8-9 repeats per concentration. Every repeat was done on ~15 worms per sex. (E) Representative confocal micrographs of a *C. remanei* female (left) and male (right) stained with the lipophilic dye DiO, showing the same sensory neurons as *C. elegans*, both in the head and in the tail. We performed a Mann-Whitney test for all comparisons, *** $P < 0.001$, ** $P < 0.01$, * $P < 0.05$, ns - non-significant.

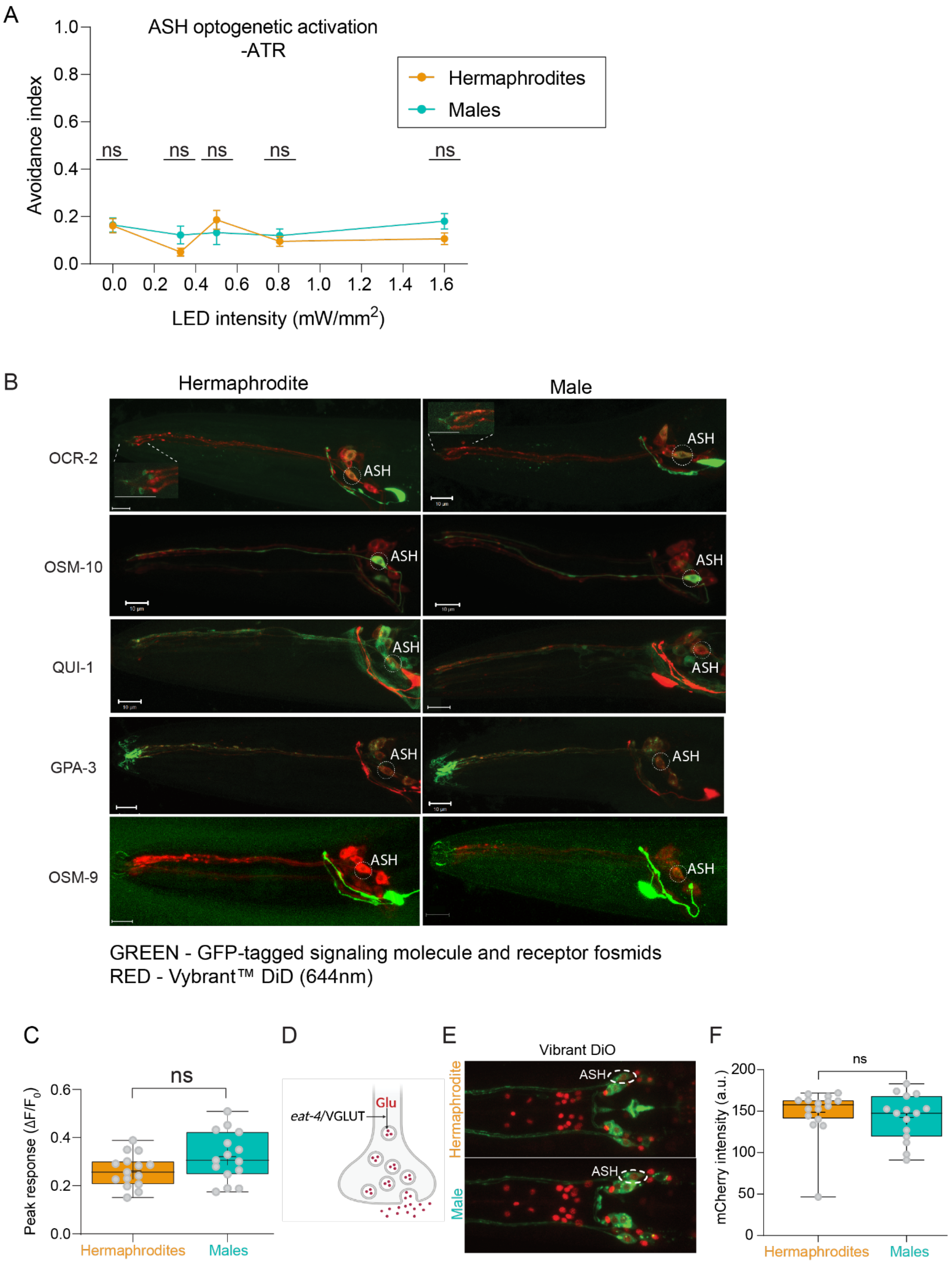


**Figure S2. ASH senses aversive stimuli similarly in the two sexes**

(A) Optogenetic activation experiments of animals without all-trans-retinal (ATR) (control groups for Figure 2D). n = 30-40 hermaphrodites, 29-38 males. (B) Representative confocal micrographs of 1 day adult hermaphrodites and males expressing fosmid reporters of candidate signaling molecules and receptor genes (*ocr-2*, *osm-10*, *qui-1*, *gpa-3*, *osm-9*) tagged with GFP, and Vybrant lipophilic dye (DiD), enabling ASH identification (marked by white dashed ovals). In *ocr-2* micrographs, the inset is a magnification of the nose, labeling subcellular expression in the dendritic tip. Some images also show the co-injection marker *ttx-3p::mCherry/GFP*. n = 14-16 animals per group. (C) Quantification of the peak responses of individual animals to ASH activity (Figure 2E-F). n = 15 hermaphrodites, 15 males. (D) Illustration of protein *eat-4*/VGLUT's function, packaging glutamate into vesicles in neurons. (E) Representative images of *eat-4*/VGLUT protein levels in ASH (marked with dashed circles), in the two sexes. ASH was identified using DiO staining (see Methods). (F) Quantification of *eat-4*/VGLUT fosmid reporter expression in ASH. n = 16 hermaphrodites, 15 males. Scale bars in all confocal micrographs are 10 µm. C and F are box and whiskers plots, vertical line represents the median and "+" is the mean. We performed a Mann-Whitney test for all comparisons, ns - non-significant.

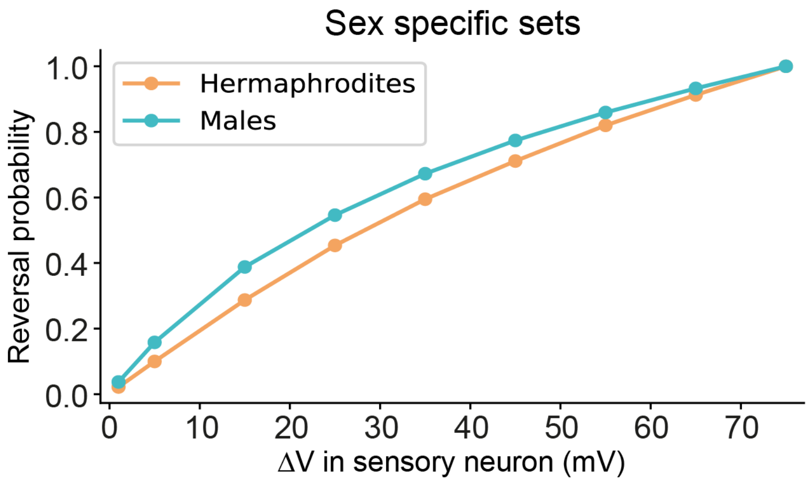


**Figure S3. Networks using the sex-specific sets for each sex do not reproduce the dimorphic response**

X axis – voltage increase in the sensory neuron following the stimulus (see Methods). Y axis – reverse probability averaged over all sets (15,992 for the hermaphrodites, 9,537 for the males). The difference in the activation of motor neurons A and B was taken as the indicator for the movement's direction.

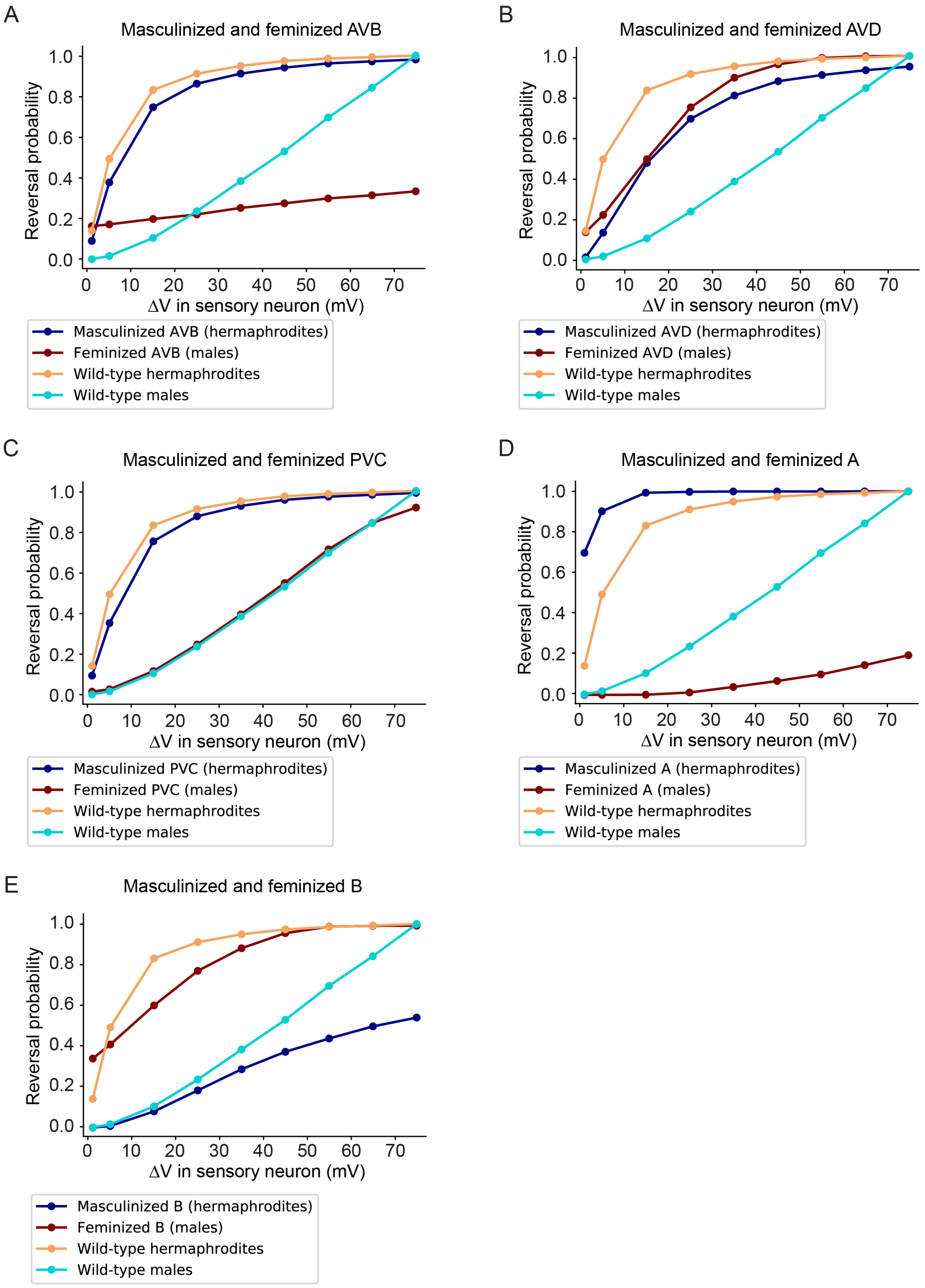
A
Masculinized and feminized AVB
Reversal probability
ΔV in sensory neuron (mV)
Masculinized AVB (hermaphrodites)
Feminized AVB (males)
Wild-type hermaphrodites
Wild-type males
B
Masculinized and feminized AVD
Reversal probability
ΔV in sensory neuron (mV)
Masculinized AVD (hermaphrodites)
Feminized AVD (males)
Wild-type hermaphrodites
Wild-type males
C
Masculinized and feminized PVC
Reversal probability
ΔV in sensory neuron (mV)
Masculinized PVC (hermaphrodites)
Feminized PVC (males)
Wild-type hermaphrodites
Wild-type males
D
Masculinized and feminized A
Reversal probability
ΔV in sensory neuron (mV)
Masculinized A (hermaphrodites)
Feminized A (males)
Wild-type hermaphrodites
Wild-type males
E
Masculinized and feminized B
Reversal probability
ΔV in sensory neuron (mV)
Masculinized B (hermaphrodites)
Feminized B (males)
Wild-type hermaphrodites
Wild-type males

**Figure S4. Masculinizing and feminizing cells of the circuit have different behavioral effects** (A-E) Computational model predictions of backward movement in wild-type animals and animals with one feminized or masculinized cell. X axis – voltage increase in the sensory neuron following the stimulus (see Methods). Y axis – reverse probability averaged over the intersection sets of both sexes. (A) Feminized and masculinized AVB neurons (all the dimorphic connections of AVB are sex-switched). Backward movement is predicted to decrease in feminized males below that typical in males. In masculinized hermaphrodites, the avoidance is not significantly affected. (B) Feminized and masculinized AVD neurons. Backward movement is predicted to increase in feminized males and insignificantly decrease in masculinized hermaphrodites. (C) Feminized and masculinized PVC neurons. Switching the connections of PVC did not result in a behavioral change for either sex. (D) Feminized and masculinized A neurons. Backward movement is predicted to decrease in feminized males below that typical in males. Accordingly, in masculinized hermaphrodites, the avoidance slightly increased. (E) Feminized and masculinized B neurons. Backward movement is predicted to significantly increase in feminized males and significantly decrease in masculinized hermaphrodites.

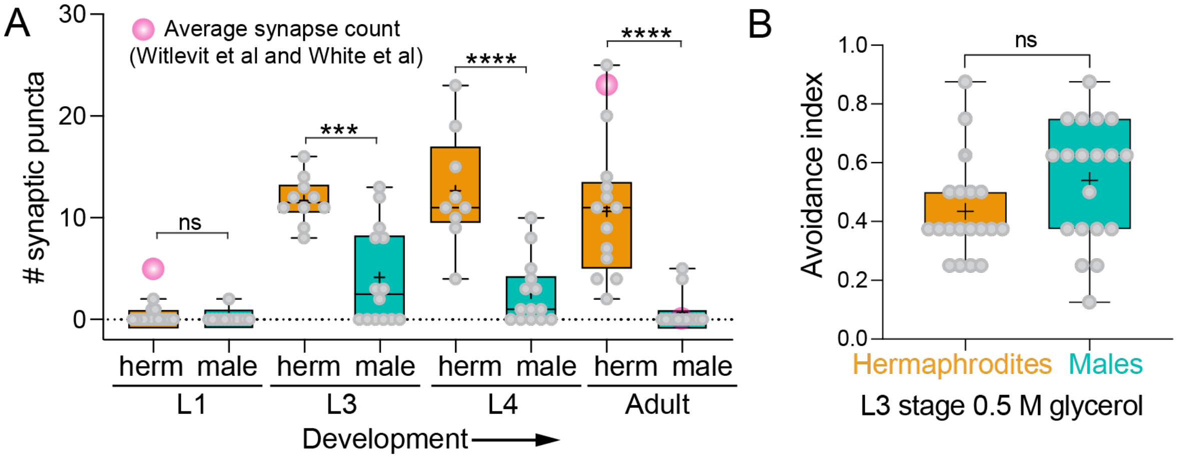


**Figure S5. Quantification of ASH-AVA iBLINC GFP puncta in hermaphrodites and males at four developmental stages.**

(A) n = 9-15 for each group. Pink circles represent the average synapse count from electron microscopy reconstructions for hermaphrodites (White et al., 1986; Witvliet et al., 2021) and males (Cook et al., 2019). Male electron-microscopy data exists only for the adult. (B) Quantification of the behavioral responses of juvenile L3 animals to aversive stimuli. The avoidance index in the tail-drop assay is calculated as in Figure 1. All plots are box and whiskers plots, vertical line represents the median and "+" is the mean. We performed a Mann-Whitney test for all comparisons, **** $P < 0.0001$, *** $P < 0.001$, ns - non-significant.

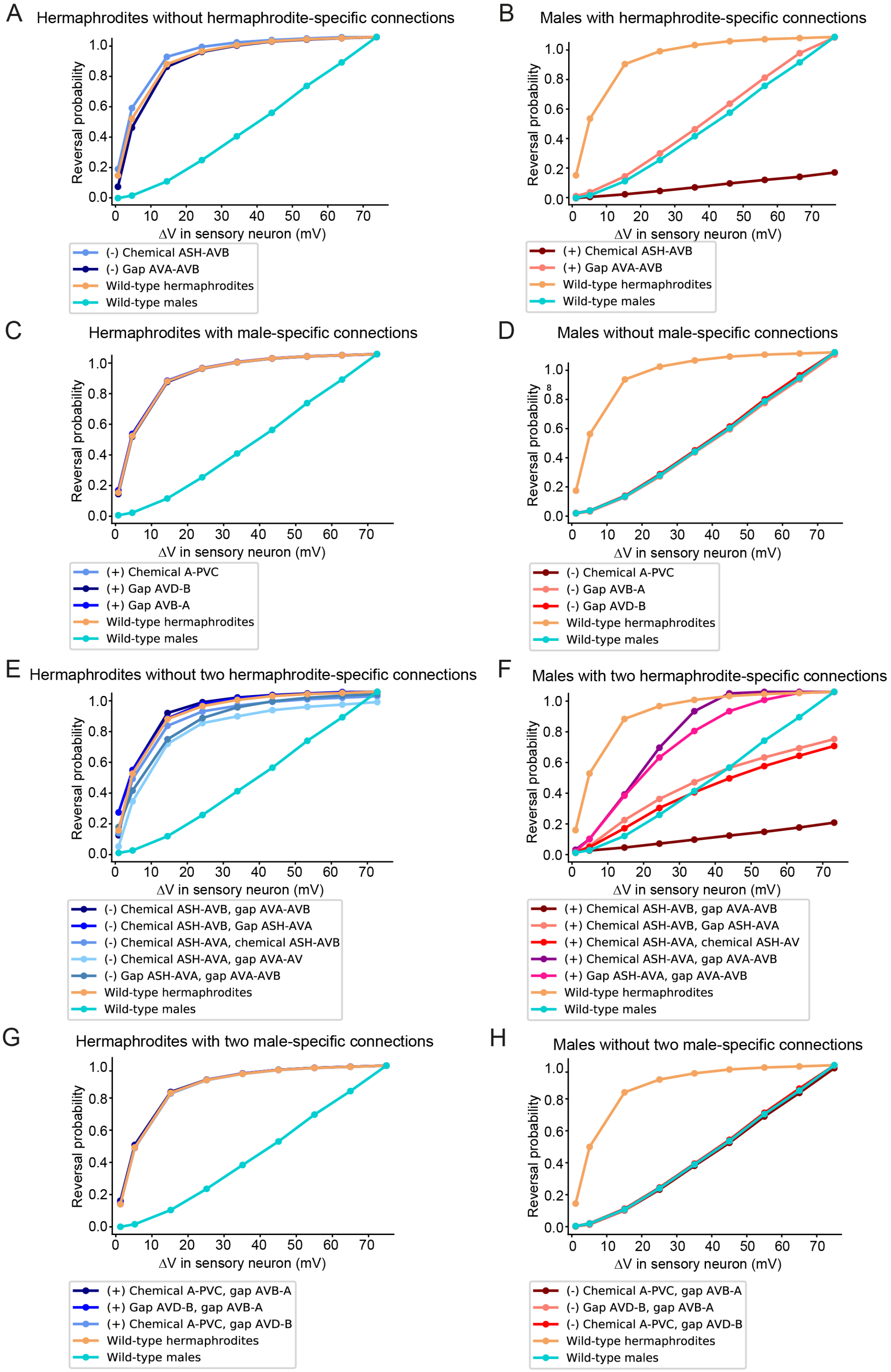
A
Hermaphrodites without hermaphrodite-specific connections
Reversal probability
ΔV in sensory neuron (mV)
(-) Chemical ASH-AVB
(-) Gap AVA-AVB
Wild-type hermaphrodites
Wild-type males
B
Males with hermaphrodite-specific connections
Reversal probability
ΔV in sensory neuron (mV)
(+) Chemical ASH-AVB
(+) Gap AVA-AVB
Wild-type hermaphrodites
Wild-type males
C
Hermaphrodites with male-specific connections
Reversal probability
ΔV in sensory neuron (mV)
(+) Chemical A-PVC
(+) Gap AVD-B
(+) Gap AVB-A
Wild-type hermaphrodites
Wild-type males
D
Males without male-specific connections
Reversal probability
ΔV in sensory neuron (mV)
(-) Chemical A-PVC
(-) Gap AVB-A
(-) Gap AVD-B
Wild-type hermaphrodites
Wild-type males
E
Hermaphrodites without two hermaphrodite-specific connections
Reversal probability
ΔV in sensory neuron (mV)
(-) Chemical ASH-AVB, gap AVA-AVB
(-) Chemical ASH-AVB, Gap ASH-AVA
(-) Chemical ASH-AVA, chemical ASH-AVB
(-) Chemical ASH-AVA, gap AVA-AV
(-) Gap ASH-AVA, gap AVA-AVB
Wild-type hermaphrodites
Wild-type males
F
Males with two hermaphrodite-specific connections
Reversal probability
ΔV in sensory neuron (mV)
(+) Chemical ASH-AVB, gap AVA-AVB
(+) Chemical ASH-AVB, Gap ASH-AVA
(+) Chemical ASH-AVA, chemical ASH-AV
(+) Chemical ASH-AVA, gap AVA-AVB
(+) Gap ASH-AVA, gap AVA-AVB
Wild-type hermaphrodites
Wild-type males
G
Hermaphrodites with two male-specific connections
Reversal probability
ΔV in sensory neuron (mV)
(+) Chemical A-PVC, gap AVB-A
(+) Gap AVD-B, gap AVB-A
(+) Chemical A-PVC, gap AVD-B
Wild-type hermaphrodites
Wild-type males
H
Males without two male-specific connections
Reversal probability
ΔV in sensory neuron (mV)
(-) Chemical A-PVC, gap AVB-A
(-) Gap AVD-B, gap AVB-A
(-) Chemical A-PVC, gap AVD-B
Wild-type hermaphrodites
Wild-type males

**Figure S6. Rewiring a single connection or two connections simultaneously had a minor effect on hermaphrodites but significantly changed male behavior**

(A-D) Computational model predictions of backward movement in wild-type animals and in animals with one connection feminized or masculinized. X axis – voltage increase in the sensory neuron following the stimulus (see Methods). Y axis – reverse probability averaged over the intersection sets of both sexes. (A) Eliminating a single hermaphrodite-specific connection in hermaphrodites did not result in a behavioral change. (B) Inserting a single hermaphrodite-specific connection into males resulted in behavioral changes. The chemical connection of ASH to AVB reduced the males' backward movement, while the gap junction between AVA to AVB didn't significantly affect their behavior. (C) In hermaphrodites, inserting a single male-specific connection did not affect their behavior. (D) In males, removing a single male-specific connection did not affect their behavior. (E-H) Computational model predictions of backward movement in wild-type animals and in animals with two connections feminized or masculinized simultaneously. (E) In hermaphrodites, removing simultaneously two hermaphrodite-specific connections resulted in a slight, insignificant decrease in their response. (F) In males, inserting simultaneously two hermaphrodite-specific connections to the males resulted in various effects. Inserting the gap junction between AVA and AVB together with either of the connections between ASH to AVA (gap junction or chemical synapse) resulted in a drastic increase in the males' avoidance response. However, inserting the gap junction between AVA and AVB together with the chemical connection between ASH and AVB resulted in a major decrease in the response. Additional combinations had only minor effects. (G) In hermaphrodites, inserting two male-specific connections did not affect their behavior. (H) In males, removing two male-specific connections did not affect their behavior.

### Key resources table

| REAGENT or RESOURCE | SOURCE | IDENTIFIER |
|---|---|---|
| Bacterial and virus strains | | |
| OP50 | Caenorhabditis Genetics Center | |
| Experimental models: Organisms/strains | | |
| *Caenorhabditis remanei* | Caenorhabditis Genetics Center | PB4641 |
| *Caenorhabditis afra* | Caenorhabditis Genetics Center | JU1286 |
| *Caenorhabditis elegans* strains: | | |
| *him-5(e1490) V* | Caenorhabditis Genetics Center | CB4088 |
| *lite-1(ce314); ljIs114[gpa-13p::FLPase, sra-6p::FTF::ChR2::YFP]* | (Ezcurra et al., 2011) | AQ2235 |
| *lite-1(ce314); ljIs114; him-5(e1490) V* | This paper | MOS52 |
| *etyIs1[gpa-13p::FLPase, sra-6p::FTF::GCaMP6s]; him-5(e1490) V*<br>6X back-crossed | This paper | MOS88 |
| *otIs518[eat-4(fosmid)::SL2::mCherry::H2B, pha-1(+)]; him-5(e1490) V* | Caenorhabditis Genetics Center | OH13645 |
| *kyIs673[sra-6p::eat-4::pHluorin, unc-122p::dsRed]* UV integrated, backcrossed to N2 8x | (Ventimiglia and Bargmann, 2017) | CX16921 |
| *kyEx4787 [rig-3p::iGluSnFR, unc-122p::dsRed]; kyEx4786 [rig-3p::RCaMP1e, unc-122p::GFP]* | (Katz et al., 2019) | OS10227 |
| *kyIs673; kyEx4786; him-5(e1490) V* | This paper | MOS198 |
| *etyEx35[flp-18p::GCaMP6s, ttx-3p::mCherry]; him-5(e1490) V* | This paper | MOS134 |

| | | |
|---|---|---|
| *akIs3[nmr-1::GFP, lin-15(+)] V* | Caenorhabditis Genetics Center | VM484 |
| *akIs3; him-8(e1489) IV* | This paper | MOS261 |
| *etyEx70[pMO34(flp-18p::tra-2[IC]), unc-122p::gfp]; lite-1(ce314); ljIs114; him-5(e1490) V* | This paper | MOS279 |
| *etyEx71[pMO30 (flp-18p::fem-3::SL2::2XNLS::tagRFP), unc-122p::gfp]; lite-1(ce314); ljIs114; him-5(e1490) V* | This paper | MOS280 |
| *him-5(e1490)V; fsEx357[osm-5p::TRA-2ic::mCherry]* | (Fagan et al., 2018) | UR754 |
| *fsEx357; lite-1(ce314); ljIs114; him-5(e1490) V* | This paper | MOS369 |
| *him-5(e1490)V; fsIs22[osm-5p::fem-3::mCherry, unc-122p::GFP]* (LG unknown) | This paper | UR1094 |
| f*sIs22; lite-1(ce314); ljIs114; him-5(e1490) V* | This paper | MOS370 |
| *etyEx106[sra-6p::Cx36::YFP, flp-18p::Cx36::mCherry; unc-122p::GFP]; him-5(e1490) V* | This paper | MOS374 |
| N2: Ancestral from CGC | Caenorhabditis Genetics Center | WormBase ID: WBStrain00000003 |
| *etyEx22[ocr-2::GFP fosmid, ttx-3p::mCherry]; him-5(e1490) V* | This paper | MOS100 |
| *nuIs11[osm-10::GFP, lin-15(+)]* | Caenorhabditis Genetics Center | HA3 |
| *nuIs11; him-5(e1490) V* | This paper | MOS69 |
| *etyEx86[qui-1::GFP fosmid, ttx-3p::mCherry]; him-5(e1490) V* | This paper | MOS334 |
| *etyEx109[gpa-3::GFP fosmid, ttx-3p::mCherry]; him-5(e1490) V* | This paper | MOS380 |
| *etyEx42[osm-9::GFP fosmid, ttx-3p::mCherry]; him-5(e1490) V* | This paper | MOS192 |
| *etyEx121[sra-6p::BirA::nrx-1, pMO26 (flp-18p::AP::NLG-1), unc-122::streptavidin::2xsfGFP, MVC11 (flp-18p::mcherry), pRF4(rol-6)]; him-5(e1490)* | This paper | MOS435 |

| | | |
|---|---|---|
| *etyEx71; etyEx121; him-5(e1490)* | This paper | MOS441 |
| *fsIs22; etyEx121; him-5(e1490)* | This paper | MOS464 |
| *fsEx357; etyEx121; him-5(e1490)* | This paper | MOS469 |
| *etyEx70; etyEx121* | This paper | MOS474 |
| *unc-31(e928) IV* | Caenorhabditis Genetics Center | DA509 |
| Oligonucleotides | | |
| CCCAGCTTTCTTGTACAAAGTGGGAATGGTTGATTCAAGTAGAAGA | ASH-specific GCaMP6s | GCaMP3_F1 |
| CACCATGGTGGCGGCCGCGGGTTTAGCCGTCATCATCTGAACG | | GCaMP3_R1 |
| CGTTCAGATGATGACGGCTAAACCCGCGGCCGCCACCATGGTG | | sra-6bb_F1 |
| TCTTCTACTTGAATCAACCATTCCCACTTTGTACAAGAAAGC | | sra-6bb_R1 |
| TAAAGAATTCCAACTGAGCGC | Masculinization and feminization of ASH | pTNZ109 F3u |
| TCCCACTTTGTACAAGAAAGCTG | | pTNZ109 R3u |
| CAGCTTTCTTGTACAAAGTGGGAatggaattctcaatcaaacgatc | | tra-2IC_SL2tagRFP_F1 |
| GCGCTCAGTTGGAATTCTTTAtcagttggaattcgaagcttg | | tra-2IC_SL2tagRFP_R1 |
| CAGCTTTCTTGTACAAAGTGGGAatggaggtggatccgggttca | | fem-3_SL2tagRFP_F1 |
| AACTTTGTATAGAAAAGTTGAAATCTGTCACATACTGCTCGAATCG | AVA-specific Cx36 construct | MVC12_F |

| | | |
|---|---|---|
| GCTTTTTTGTACAAACTTGTTTCGGGGGTAGATTTCAAATAGATTGG | | MVC12_R |
| ATTTGAAATCTACCCCCGAAACAAGTTTGTACAAAAAAGCAG | | pIR204_Fwd |
| GAGCAGTATGTGACAGATTTCAACTTTTCTATACAAAGTTGATAGC | | pIR204_Rev |
| ttattatttcagattttgccGGATCCCCGGGATTGGCCAA | ASH-specific iBlinc construct | pMO22_Fwd |
| cgatttattatatctaaaagATTTCATTTCCAAGTTGTTAGCGTATCC | | pMO22_Rev |
| CTAACAACTTGGAAATGAAATcttttagatataataaatcgaaattgaaatgt | | Psra-6_Fwd |
| TTGGCCAATCCCGGGGATCCggcaaaatctgaaataataaatattaaattc | | Psra-6_Rev |
| ggaggacccttggctagcATGGTTGATTCAAGTAGAAGAAAATG | AVA-specific GCaMP6s | Lada7F |
| ggcgctcagttggaattctTTAGCCGTCATCATCTGAAC | | Lada7R |
| GCTAGCCAAGGGTCCTCCTGAA | | GRASPbb _R1u |
| GAATTCCAACTGAGCGCCGGTCG | | GRASPbb_F1u |
| Software and algorithms | | |
| MATLAB | | https://www.mathworks.com/products/matlab.html |
| FIJI | | https://imagej.net/software/fiji/ |
| BioRender | | https://biorender.com/ |

| | | |
|---|---|---|
| GraphPad Prism | | https://www.graphpad.com/scientific-software/prism/ |
| Adobe Illustrator | | https://www.adobe.com |
| WormLab | | https://www.mbfbioscience.com/wormlab |
| ZEN | | https://www.zeiss.com/microscopy/int/products/microscope-software/zen.html |
| Python - Spyder | | https://www.spyder-ide.org/ |
